# Rhizosphere Bacteria and Fungi are Differentially Structured by Host Plants, Soil Mineralogy and Ectomycorrhizal Communities in the Alaskan Tundra

**DOI:** 10.1101/2025.01.29.635592

**Authors:** Sean Robert Schaefer, Fernando Montaño-López, Hannah Holland-Moritz, Caitlin Hicks Pries, Jessica Gilman Ernakovich

## Abstract

The rhizosphere contains a diverse group of bacteria and fungi living near plant roots whose composition and function are key drivers of ecosystem and biogeochemical processes. Despite rich literature on rhizosphere communities, surprisingly few studies have examined the drivers of rhizosphere community structures in natural settings. We collected 513 root samples from 141 individual plants representing six plant species and three mycorrhizal association types across four glacial drifts in the North Slope of Alaska. Glacial drifts ranged from 11,000 to 4.5 million years since deglaciation representing a gradient in glacial history and mineralogical weathering. We found that glacial history, a strong proxy for soil mineralogy, explained most of the captured variation in rhizosphere bacterial communities (13.3%) and ectomycorrhizal fungal communities (10.2%) while interactions between glacial history and host plants explained the most variation in fungal rhizosphere communities (11.6%). We analyzed ectomycorrhizal fungal communities from the shrub *Betula nana* across spatial scales and sites and found a large correlation between ectomycorrhizal and rhizosphere communities, and that ectomycorrhizal composition was most similar among root fragments belonging to the same plant, followed by plants at the same site, and were most dissimilar for plants at different sites.

## 1. Introduction

The rhizosphere describes the area directly surrounding a root while mycorrhizosphere surrounding a mycorrhizal hyphae (Hartmann et al., 2009, van Elsas and Boersma 2011). This interface between roots and soil or hyphae and soil is critical to plant health, soil ecology, and biogeochemistry, as it is the primary site for C and nutrient exchange (Jones et al., 2009). An estimated 5-40% of photosynthates formed through primary production are secreted as exudates from roots and hyphae to the surrounding soil making the rhizosphere a primary location for transferring atmospheric C to belowground pools (Nguyen 2003, Badri and Vivanco 2009, Dennis et al., 2010, el Zahar Haichar et al., 2014). Rhizosphere bacterial and fungal community compositions are a product of plant selection through exudation, environmental selection through soil edaphic factors (Jones et al., 2019), and random or stochastic processes (Liu et al., 2023). Plants recruit microbes to the rhizosphere by altering the magnitude, timing, and composition of their root exudate profiles, enabling some microbes to grow and colonize while inhibiting the growth of others (Dennis et al., 2010). The tundra is a unique model for studying the rhizosphere communities, as global change is causing large shifts in vegetation communities. Regional-scale displacement of non-mycorrhizal sedges and mosses by mycorrhizal, woody shrubs are rapidly occurring in a process called shrubification (Sturm et al., 2001, Myers-Smith et al., 2011, Heijmans et al., 2022). Common tundra plants impacted by shrubification include various functional types such as graminoids (e.g., *Eriophorum vaginatum*), deciduous shrubs (e.g., *Betula nana, Salix pulchra, Salix reticulata.*), evergreen shrubs (e.g., *Empetrum nigrum*), heath (e.g., *Vaccinium uliginosum*), lichens, and mosses (Wullschleger et al., 2014). These functional types differ in belowground traits including root architecture, rooting depth, root exudate profiles, and mycorrhizal association (Iversen et al., 2015). For example, evergreen and heath shrubs are obligate ericoid mycorrhizal (ERI) (Leopold 2016), whereas graminoids are primarily non-mycorrhizal (Chapin and Kӧrner 1995), although can form rare relationships with arbuscular mycorrhizae in dry habitats (Muthukumar et al., 2004). Deciduous shrubs are obligate ectomycorrhizal (ECM) (Miller 1982, Treu et al., 1996, Ryberg et al., 2009) and represent the largest “winners” of shrubification as exemplified by their substantial increases in prevalence and biomass, as well as expanded northward range in ice-poor permafrost regions (Mekonnen et al., 2021, Heijmans et al., 2022). ECM fungi can have strong effects on fungal and bacterial saprotrophic (decomposer) activity and composition (Gadgil and Gadgil 1971, Lindahl et al., 2007, Kluber et al., 2011, Bӧdeker et al., 2016, Fernandez and Kennedy 2016, Shirakawa et al., 2019, Fitch et al., 2020, Fitch et al., 2023). Thus, the increase in ECM-associated shrubs is likely to alter microbial communities and may produce perturbations in biogeochemical processes, such as the carbon and nitrogen cycles (Parker et al., 2021).

Soil mineralogy plays a key role in shaping the underlying physical and chemical environments that influence microbial composition in the bulk soil and rhizosphere (Ahmed et al., 2017, Finley et al., 2018, Jilling et al., 2018, Whitman et al., 2018, Finley et al., 2021). In the tundra, glacial retreat, parent material, and the length of time minerals have been exposed to weathering control soil mineralogy (Whittinghill & Hobbie 2011). As glaciers move, they deposit fresh bedrock onto the soil and significant weathering only begins when the glaciers retreat (Southard 2022). The time since the last deglaciation, as well as the parent material being deposited, are strong proxies for the stage of mineralogical transformation and weathering (Munroe and Bockheim 2001). Differences in the soil’s physical and chemical environments brought on by differences in mineralogy result in distinct microbial communities in the bulk soil (Bakke and Ernakovich, unpublished; Ahmed et al., 2017; Whittinghill and Hobbie 2011). Because microbes in the rhizosphere are selected from bulk soil (deVries and Wallenstein 2017), there is a strong, indirect relationship between soil mineralogy and rhizosphere community composition; direct relationships between soil mineralogy and rhizosphere composition are complex and involve multiple factors within the plant-microbe-mineral system such as organo-mineral interactions, nutrient acquisition strategies, and edaphic factors (Finley et al., 2021). Nevertheless, there is evidence for strong associations between soil mineralogy and rhizosphere community composition (Whitman et al., 2018).

To better understand how rhizosphere communities change between different plant species, glacial histories, and ECM fungal communities, we quantified their relative importance in shaping bacterial and fungal rhizosphere communities across plants belonging to a diverse set of functional types and mycorrhizal host types along a deglaciation gradient. Our primary questions were: 1) To what degree do plant species and glacial history explain rhizosphere community composition, and is this consistent for bacterial and fungal communities? 2) What is the variation of ECM fungal composition between sites, between plants at the same site, and within a single plant? 3) To what extent are bacterial and fungal rhizosphere community compositions correlated to ECM fungal community composition, and does the strength of this correlation differ between ECM to bacterial and ECM to fungal rhizosphere communities? We hypothesized that plant species would explain the most variation in bacterial and fungal rhizosphere communities followed by glacial history. Because most ECM fungi are generalists and widespread throughout the tundra (Gardes and Dalberg 1996, Botnen et al., 2013), we hypothesized that within-site variation of ECM fungi associated with *B. nana* roots would be equal to variation between sites. We also hypothesized that the variation of ECM fungi within a single plant would be low due to interspecies ECM fungal competition for root colonization. Finally, we hypothesized that bacterial and fungal rhizosphere community compositions strongly correlate with ECM fungal communities due to mycorrhizal ability to engage in exudation and influence surrounding microbial communities (Gorka et al., 2019). This observational study deepens our understanding of how complex interactions between plant species, soil mineralogy, and mycorrhizal fungi synergistically influence the rhizosphere communities responsible for many ecosystem processes. We hope this work will serve as a foundation for better understanding the underlying microbial ecology involved in plant-microbe-mineral interactions in tundra environments.

## 2. Materials and Methods

### 2.1 Site selection and chemical analysis

Samples were collected at or near Toolik Field Station in the North Slope of Alaska (Fig. 1). Sites were chosen based on differences in their glacial history (Hamilton 2003) and plant communities (Gough et al., 2000, Greaves et al., 2019) which inferred differences in soil physico-chemical properties such as carbon, aluminum, and iron content (Sposito 2008, Whittinghill and Hobbie 2011). The vegetation communities consisted of moist acidic tundra (MAT) and moist non-acidic tundra (MNT) (Walker et al., 1994). Collection took place in August 2021 and 2022 when the active layer was approaching peak seasonal thaw depth. Land use permits for sample collection were obtained from the Bureau of Land Management (permit #F-97705) and Alaska Department of Natural Resources (written approval).

**Fig. 1.**
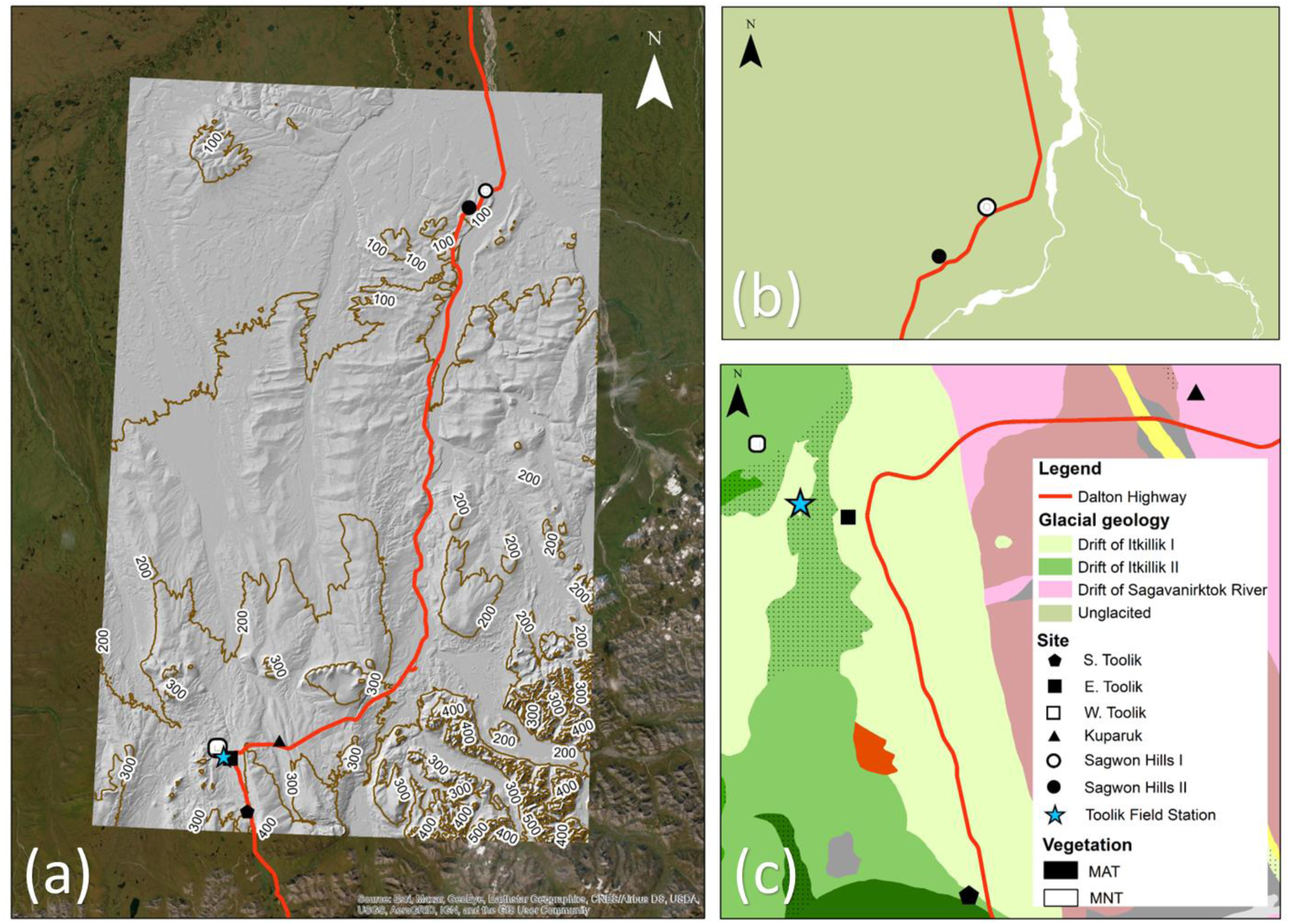
Map of sites and glacial drifts of the North Slope of Alaska near Toolik Field Station. A full topographical map of the sites **(a)** where numbers represent elevation and symbols shaded in black represent moist acidic tundra vegetation (MAT) while shaded in white is moist non-acidic tundra (MNT). Sagwon Hills sites **(b)** and Toolik sites **(c)** are displayed in higher resolution showing the different glacial drifts across the region (Hamilton 2003).

Differences in soil chemistry and mineralogy were confirmed using various chemical analyses (Table 1). Prior to chemical analysis, samples were dried and sieved (<2 mm), root-picked, and homogenized. Soil pH was measured on a Mettler Toledo S220 pH/ion meter, using a 1:2 soil-to-water ratio. Samples were finely ground prior to CN elemental analysis using Delta V advantage (Thermo Fisher Scientific, Waltham, MA). To estimate pedogenic oxides we conducted citrate dithionite extractions based on the previously described methods (Loeppert & Inskeep 1996, Carter and Gregorich 2007). Briefly, we mixed 0.5 g of finely ground soil with 6 g of sodium citrate, 0.5 g of sodium hydrosulfite, and 30 mL of Milli-Q water in 50 mL centrifuge tubes. The samples were shaken for 16 hours and centrifuged. All suspensions were filtered through Whatman 52 filter papers. The filtrates were diluted with 5% HNO_3_ to a final volume of 1:50 (v/v), and metal concentrations were measured using inductively coupled plasma-optical emission spectroscopy (ICP-OES, Spectro ARCOS).

**Table 1.** Site descriptions and analyses. Edaphic differences between sites are a product of their glacial histories. Years since deglaciation represents the approximate age since glaciers retreated and is a proxy for mineralogical weathering. Sites were given operational names based on their locations. MAT is moist acidic tundra vegetation communities; MNT is moist non-acidic tundra vegetation communities. The number of samples used in each chemical analysis is shown in subscript parentheses, aside from predominant mineral groups, where one sample was used for characterization. We did not collect soil samples at the site W. Toolik and thus did not analyze its soil chemistry or mineralogy. The values for pH, C, N, Al and Fe and predominate mineral groups are the averages of sample(s) between 0 and 15 cm depths from the surface. Mean concentrations of Al and Fe extracted by citrate-dithionite are presented along with predominate mineral groups identified with X-Ray powder diffraction analysis; OM refers to the general characterization of organic matter.

| Glacial drift | Years since de-glaciation | Site | GPS co-ordinates | Mean annual temperature (°C) | Elevation (m) | Vegetation type | pH | C (%) | N (%) | Al (mg g<sup>-1</sup>) | Fe (mg g<sup>-1</sup>) | Predominate mineral groups, (wt. %) |
| --- | --- | --- | --- | --- | --- | --- | --- | --- | --- | --- | --- | --- |
| **Itkillik II** | 11,000-25,000 | S. Toolik | 68.550<br>-149.496 | -10.5 | 893 | MAT | 5.38<br>(18) | 33.78<br>(18) | 1.03<br>(18) | 1.73 (20) | 24.9 (20) | OM, 50.8%; Quartz, 13.6%; Illite, 14.5%; Smectite, 11.5%; Chlorite, 7.3% |
| **Itkillik II** | 11,000-25,000 | W. Toolik | 68.639<br>-149.623 | -2.43 | 761 | MNT | - | - | - | - | - | - |
| **Itkillik I** | 53,000 – 66,000 | E. Toolik | 68.625<br>-149.571 | -2.43 | 761 | MAT | 4.58<br>(29) | 11.09<br>(29) | 0.38<br>(29) | 1.52 (22) | 14.6 (22) | OM, 45.5%; Quartz, 27.2%; Chlorite, 8.3%; Smectite, 7.6%; Illite 4.1% |
| **Saga-vanirktok river** | 125,000-728,000 | Ku-paruk | 68.653<br>-149.382 | -10.4 | 795 | MAT | 4.30<br>(30) | 19.03<br>(30) | 0.56<br>(30) | 1.51 (21) | 13.4 (21) | OM, 54.1%; Quartz, 23.5%; Smectite, 8.0%; Chlorite, 6.3%; Illite, 2.8% |
| **Unglaciaded** | 4,500,000 | Sag-won Hills I | 69.450<br>-148.629 | -11.7 | 214 | MNT | 5.92<br>(6) | 31.14<br>(6) | 1.28<br>(6) | 0.99 (1) | 9.38 (1) | OM, 43.1%; Quartz, 29.8%; Chlorite, 8.73%; Plagioclase, 6.2%; Vermiculite, 3.6%; |
| **Unglaciaded** | 4,500,000 | Sag-won Hills II | 69.425<br>-148.694 | -11.7 | 214 | MAT | 5.34<br>(5) | 7.85<br>(5) | 0.44<br>(5) | 1.32 (2) | 9.80 (2) | - |

Identification of crystalline minerals as well as organic matter components were performed through X-ray powder diffraction (XRPD). In brief, finely ground soil subsamples were loaded onto a Bruker D8 DISCOVER (Bruker Corporation) diffractometer equipped with a Cu-Kα radiation source. One soil sample was used for characterization and repeated to ensure diffractograms and phases matched. Diffraction pattern peak matching and quantification of crystalline minerals and organic components from the XRPD data was conducted using the automated full pattern summation (afps) function implemented in the powdR package v1.3.0 for R (Butler & Hillier 2021) and is further discussed in von Fromm et al. (2024).

### 2.2 Plant selection

Six plant species were chosen based on their abundance in the Alaskan tundra and representation of functional types (Walker et al., 1994, Treu et al., 1995). We chose woody shrubs and ECM fungal hosts *Betula nana* (Molina et al., 1992), *Salix reticulata* (Treu et al., 1996, Ryberg et al., 2009), and *Salix pulchra* (Miller 1982); woody heath and ERI fungal hosts *Empetrum nigrum* (Treu et al., 1996, Koizumi and Nara 2017) and *Vaccinium uliginosum* (Yang et al., 2020); and the predominately non-mycorrhizal sedge, *Eriophorum vaginatum* (Chapin and Kӧrner 1995, Parker et al., 2015). Although plants in the *Salix* genera can form arbuscular, ectomycorrhizal, and even ericoid symbioses based on the species and environment (Cazares 1992), to our knowledge, *S. reticulata* and *S. pulchra* have only been reported as ECM fungal hosts in the Alaskan tundra (Miller 1982, Treu et al., 1995, Ryberg et al., 2009).

### 2.3 Sample collection

Plant roots were collected from the first 10 cm of the organic soil horizon. Entire plants were dug out and roots were separated using a Hori Hori knife and abscised using wire cutters. All instruments were sterilized with 70% ethanol prior to sampling. Roots belonging to the same plant were separated into 5-10 cm long fragments and both root fragments and soil that tightly clung to them were retained for downstream DNA extraction. Because our second question focused on variation across scales from within-plant to across sites, each root fragment was considered an individual microbial community (even if the roots came from the same plant). We collected five root sub-samples in 2021 and three sub-samples in 2022 as bacterial and fungal rhizosphere species richness was adequately represented with three samples.

### 2.4 Rhizosphere and ectomycorrhizal DNA extraction

Rhizosphere communities were obtained by washing plant root fragments in a 0.9% NaCl solution for 15 minutes (Barrilot et al. 2013). The “rhizosphere” community included both those microorganisms tightly bound to the roots (rhizoplane), and those present in the soil that clung to the roots (rhizosphere). Root fragments, approximately 0.66 grams in weight and 5cm long, were placed in a 15 ml falcon tube containing 10 ml of saline solution and vigorously shaken on a vortex adapter for 15 minutes and then centrifuged at 10,000 g for 15 minutes. The supernatant was removed, and 2 ml of organic slurry was transferred to a microcentrifuge tube where the samples were again centrifuged at 13,000 g for 2 minutes to form a pellet. The supernatant was again removed leaving approximately 0.5 – 1.0 ml of organic slurry which was then used as the starting material for DNA extraction following the Qiagen Powersoil Pro Kit protocol (Qiagen, Hilden, Germany). ECM fungi were extracted from washed *B. nana* root*s* and ground using a mortar and pestle. DNA from the ground plant roots was extracted using Qiagen Plant Mini Kit (Qiagen, Hildenm Germany). In total, we collected and processed 94 *B. nana* root samples.

### 2.5 DNA amplification and sequencing

Rhizosphere DNA was amplified using 16S rRNA and ITS region primer sets and ECM fungi with the ITS set. The 16S primers were 515F/ 926R (Parada et al., 2016) while ITS were ITS4-FUN/ 5.8S-FUN (Taylor et al., 2016). Primers contained either Nextera (16S) or TruSeq (ITS) adapters to allow for use in the Novaseq platform. PCR reaction volumes were carried out with 20-200 ng of template DNA in 12 µL. The final concentration of the reaction was 1X DreamTaq master mix (ThermoScientific, Waltham, MA), 0.3 µmol/µL forward and reverse primers and 0.05 µg/µL BSA. Bovine serum albumin (BSA; 0.25 µg/µL) replaced PCR-grade water in our PCR reactions to reduce PCR inhibition and increase DNA amplification. Our thermocycler settings for all 16S amplification were an initial 95 °C start for 2.5 minutes, followed by cycles of 94 °C for 45 seconds, 50 °C for 1 minute and 72 °C for 1.5 minutes, and was repeated for 35 cycles. Our thermocycler settings for all ITS amplification was an initial 95 °C start for 2 minutes, followed by cycles of 94 °C for 30 seconds, 58 °C for 40 seconds and 72 °C for 2 minutes, and was repeated for 35 cycles. In both cases, a 72 °C annealing step for 10 minutes was done followed by a 4 °C infinite hold. Sequencing was performed at the Hubbard Genome Center at the University of New Hampshire using paired end reads at 250 bp on an Illumina Novaseq platform.

### 2.6 DNA processing and generation of species matrices

Average sequence depths for rhizosphere bacteria and fungi were 79,125 and 125,570 respectively and ranged between 5,194 - 3,095,927 reads for bacteria and 5,124 – 1,771,462 reads for fungi. Average sequence depth for ECM fungi was 15,581 and ranged between 103 – 66,572. Sequences were processed, filtered, and trimmed using DADA2 v.1.16 (Callahan et al., 2016). Minor modifications to the pipeline were made to optimize for Novaseq sequencing platforms (Teixeira et al., 2024); this included removing any reads with 20 or more polyG’s and implementing an error rate learning step that allowed for more accurate estimation of error rates from binned quality scores, an artifact of Novaseq sequencing platforms. All primers and polyG sequenced were removed with Cutadapt v.3.5. 16S sequences were then filtered to remove low-quality reads. ITS sequences were filtered but not trimmed due to the variable lengths of the PCR product. Instead, we used ITSxpress to identify and extract ITS regions (Rivers et al., 2018). All 16S and ITS sequences were denoised using DADA2 software. Taxonomy was assigned from amplicon sequence variants (ASVs) using Naïve Bayes classifiers. The databases used were SILVA v.138.1 for bacteria and UNITE v.8.3 for fungi. In total, we found 283,736 bacterial and 35,296 unique fungal ASVs. We did not find any archaeal ASVs. Our post-DADA2 processing, including normalization to median sequencing depth (McMurdie 2018) and removal of chloroplast and mitochondrial DNA, was done in R using the phyloseq package (v1.42.0; McMurdie and Holmes 2013). Samples with less than 5,000 reads were removed from rhizosphere matrices and less than 100 reads were removed from ECM matrices. The *Betula*-ECM fungal species matrix was filtered to contain only known ectomycorrhizal species using the Fungal Traits database (Põlme et al., 2020) and a total of 81 samples with 1,467 ECM fungi ASVs remained.

### 2.7 Ordinations, Plots, and Statistical Analyses

All visualizations and statistical analyses were performed using R Statistical Software (v4.2.3; R Core Team 2023). To compare the number of unique and shared taxa across plants we created upset plots using the upset_pq function in MiscMetabar (v.0.9.2; Taudière 2024). To visualize differences in microbial composition, we created ordinations using the phyloseq package (v1.42.0; McMurdie and Holmes 2013) in conjunction with ggplot2 (v.3.4.3; Wickam 2016). Supplemental organization on phyloseq objects was done using the R package phylosmith (v1.0.7; Smith-Schuyler 2023). Heat plots were made to visualize common taxa across glacial drifts using the pheatmap function in the pheatmap package (v. 1.0.12; Kolde 2019). To quantify and compare variation in rhizosphere communities associated with various factors, we conducted permutational multivariate analysis of variance (PERMANOVA) tests in R using the adonis2 function in the vegan package (v2.6-4; Oksanen et al., 2022). For bacterial and fungal rhizosphere PERMANOVA tests, we removed all samples from W. Toolik, due to unequal representation from host plants. The total samples after removing low read samples and unbalanced sites were 340/396 rhizosphere bacteria, and 301/384 rhizosphere fungi (Table S1). Bray-Curtis dissimilarity distances were used for our species matrices due to the high number of zeros in our dataset. Our bacterial and fungal rhizosphere model was: distance of species matrix ∼ glacial history*host plant species + vegetation type; strata = year, permutations = 999 (Table 2). We dropped the factors mycorrhizal type and site from the rhizosphere models because they were confounded with host plant and glacial history respectively. However, those factors were also statistically significant (Table S2). Ultimately, we chose the models that explained the most amount of variation while minimizing confounding variables. To examine the influence of mycorrhizal type on rhizosphere composition we conducted additional PERMANOVA’s for bacteria and fungi using the model: distance of species matrix ∼ glacial history + mycorrhizal type + vegetation type (Table S2). To examine the effect of glacial history on ECM composition we used the model: distance of species matrix ∼ glacial history + vegetation type + individual plant (strata = year, permutations = 999). To examine the between site effects on ECM composition we used the model: distance of species matrix ∼ site + vegetation type + individual plant (Table 2, S2). For within-site ECM fungal subsets we used the model: distance of species matrix ∼ individual plant (Table S3).

**Table 2.** PERMANOVA results from rhizosphere and ECM models. P values of <0.001 indicate significant groupings for all the factors tested in the model, suggesting they explain a high degree of variance in bacterial rhizosphere communities. Glacial history refers to the most recent glacial drift and is a proxy for mineralogical weathering. Host plant refers to the plant species which rhizospheres were collected. An ‘×’ denotes an interaction effect between two variables. Vegetation type refers to either moist acidic or non-acidic plant communities. Individual plant refers to the plant which roots were subsampled from.

| Factor | Df | SumOfSqs | R2 | F | Pr(>F) |
| --- | --- | --- | --- | --- | --- |
| ***Bacterial model*** |  |  |  |  |  |
| Glacial drift | 3 | 15.02 | 0.133 | 21.21 | 0.001 |
| Host plant | 5 | 9.242 | 0.082 | 7.829 | 0.001 |
| Vegetation type | 1 | 2.702 | 0.024 | 11.45 | 0.001 |
| Glacial drift × Host plant | 15 | 11.11 | 0.099 | 3.136 | 0.001 |
| Residual | 316 | 74.61 | 0.662 | - | - |
| Total | 340 | 112.7 | 1 | - | - |
| ***Fungal rhizosphere model*** |  |  |  |  |  |
| Glacial drift | 3 | 8.310 | 0.063 | 7.906 | 0.001 |
| Host plant | 5 | 9.054 | 0.068 | 5.168 | 0.001 |
| Vegetation type | 1 | 2.595 | 0.020 | 7.405 | 0.001 |
| Glacial drift × Host plant | 15 | 15.32 | 0.116 | 2.914 | 0.001 |
| Residual | 277 | 97.05 | 0.733 | - | - |
| Total | 301 | 132.3 | 1 | - | - |
| *ECM fungal model* |  |  |  |  |  |
| Glacial history | 3 | 3.939 | 0.102 | 4.267 | 0.001 |
| Vegetation type | 1 | 1.478 | 0.038 | 4.803 | 0.001 |
| Individual plant | 19 | 15.67 | 0.406 | 2.680 | 0.001 |
| Residual | 57 | 17.54 | 0.454 | - | - |
| Total | 80 | 38.63 | 1 | - | - |

Mantel tests were conducted using the mantel function in the vegan; we chose to retain samples belonging to W. Toolik since this site was well represented within the *B. nana* subset (Table S1). This resulted in 52 samples for rhizosphere bacteria to ECM fungi comparison and 40 samples for rhizosphere fungi to ECM fungi. Bray-Curtis dissimilarity distance matrices were created from our trimmed species matrices and ran using Pearson correlations. Differential abundance analyses were conducted in DESeq2 in R (v1.44.0; Love et al., 2014). To examine the differences in functional potential for rhizosphere bacteria across host plants and glacial drifts, we used the FAPROTAX database and compared the average number of ASVs for each functional annotation found across host plants or glacial drifts (Louca et al., 2016).

## 3. Results

### 3.1 Rhizosphere communities are conserved independently of glacial history, but plant host type determines a unique composition of core rhizosphere taxa

Regardless of the host plant and glacial history, we found that a handful of bacterial and fungal taxa dominate most rhizosphere communities. The families with the highest relative ASV abundance for bacterial rhizosphere communities were *Acidobacteriaceae* (Subgroup 1), *Xanthobacteraceae, Acetobacteraceae, Isosphaeraceae* and *Chitinophagaceae*; these ranged from 8.5% to 5.6% relative abundance. The families with the highest relative abundance for fungal rhizosphere communities were *Hyaloscyphaceae, Herpotrichiellaceae, Mortierellaceae, Serendipitaceae*, and *Helotiaceae;* these ranged from 11.4% to 5.1% relative abundance (SI Relative Abundance dataset). We found that the 15 most abundant bacterial genera comprised 50% of the relative abundance of bacterial ASVs, and the 15 most abundant fungal genera comprised 56% of the relative abundance of fungal ASVs (SI Relative Abundance dataset). The vast majority of bacterial functional types, based on FAPROTAX, were predicted to be heterotrophs. Differences in the relative abundance of functional types across host plants (Fig. S1) and glacial drifts (Fig. S2) were minimal. However, notable differences include a decreased relative abundance of oxygenic phototrophy and photosynthetic cyanobacteria annotations in *E. vaginatum* microbiomes and a reduction of dark hydrogen oxidation and reductive acetogenesis annotations in *E. nigrum*. Likewise, increased relative abundance of aliphatic non-methane hydrocarbon degradation and aromatic hydrocarbon degradation annotations were found in *E. nigrum* relative to other host plants (Fig. S1). Notable differences in the relative abundance of functional types across glacial drifts include a reduced relative abundance of cellulolysis and manganese oxidation in unglaciated soils, and the absence of any ASVs annotated for dark oxidation of sulfur compounds in Sagavanirktok river glacial drift (Fig. S2).

Overall, there was higher bacterial rhizosphere alpha diversity than fungal. This ranged from a Shannon index between 4 and 7.5 for bacteria and 1 and 5.5 for fungi (Fig. S3). Differential abundance analyses revealed that 782 bacterial taxa and 1069 fungal taxa were significantly enriched when comparing ASVs found across different plant types (Fig. S4, SI Differential Abundance dataset), while 2882 bacteria and 769 fungi were enriched when comparing across glacial history (Fig. S5, SI Differential Abundance dataset). After removing ASVs appearing less than 10 times in the dataset; we found approximately 12.7% of bacterial ASVs and 5.5% of fungal ASVs were present across all tundra plants, while 4.2% - 8.1% of bacterial ASVs were unique to one plant species, and 3.3% - 17.8% of fungal ASVs were unique to one plant species (Fig. 2). By using Upset plots, we offer a unique way to compare the number of common features, such as ASVs, that are shared across disparate factors, such as host plants.

**Fig. 2.**
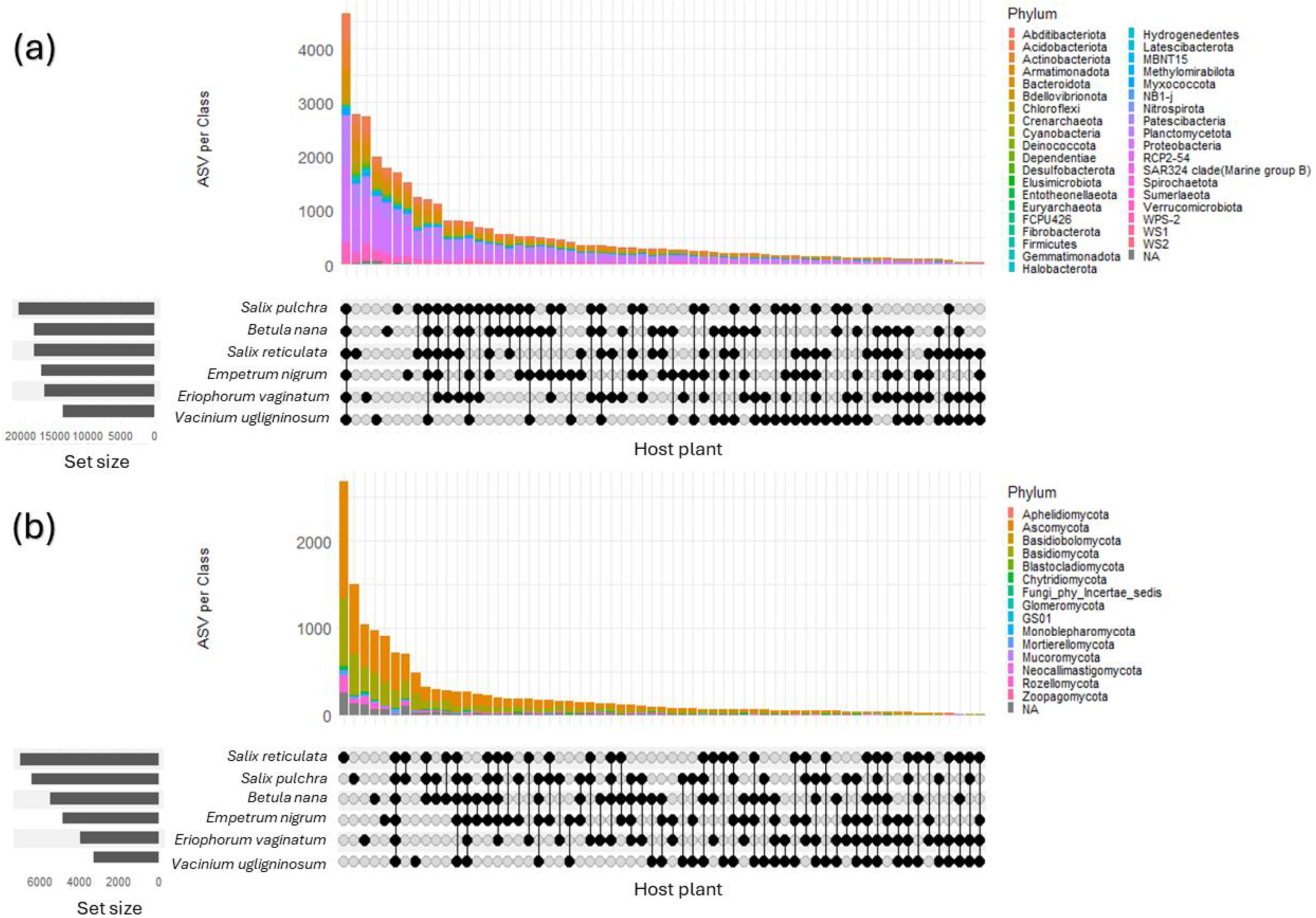
Host plant species contain a core microbiome shared across tundra plants while also displaying unique cultivation of bacteria (a) and fungi (b) for each host plant. Upset plots show the number of ASVs shared between various host plants broken down by phyla. Filled circles indicate the host plant(s) included in each comparison (class), while the set size represents the total number of ASVs belonging to each host plant. ASVs containing less than 10 reads in the dataset were removed from the plots for a total of 36,315 bacterial ASVs and 14,684 fungal ASVs.

### 3.2 Glacial history and host plants are among the strongest factors that structure bacterial and fungal rhizosphere communities

Glacial history, host plant species, and the interaction of glacial history and host plant species shape both the bacterial and fungal composition (Table 2). Between a quarter to a third of the variation in bacterial (31.4%) (Fig. 3a, Table 2) and fungal (24.7%) rhizospheres (Fig. 3b, Table 2) were captured in our models using the factors glacial history, host plant, and their interaction. The explanatory power for host plant was nearly equal between bacterial and fungal communities at 8.2% and 6.8%, respectively. Glacial history however had a larger gap between the two groups of microorganisms at 13.3% and 6.3% for bacteria than fungi, respectively

**Fig. 3.**
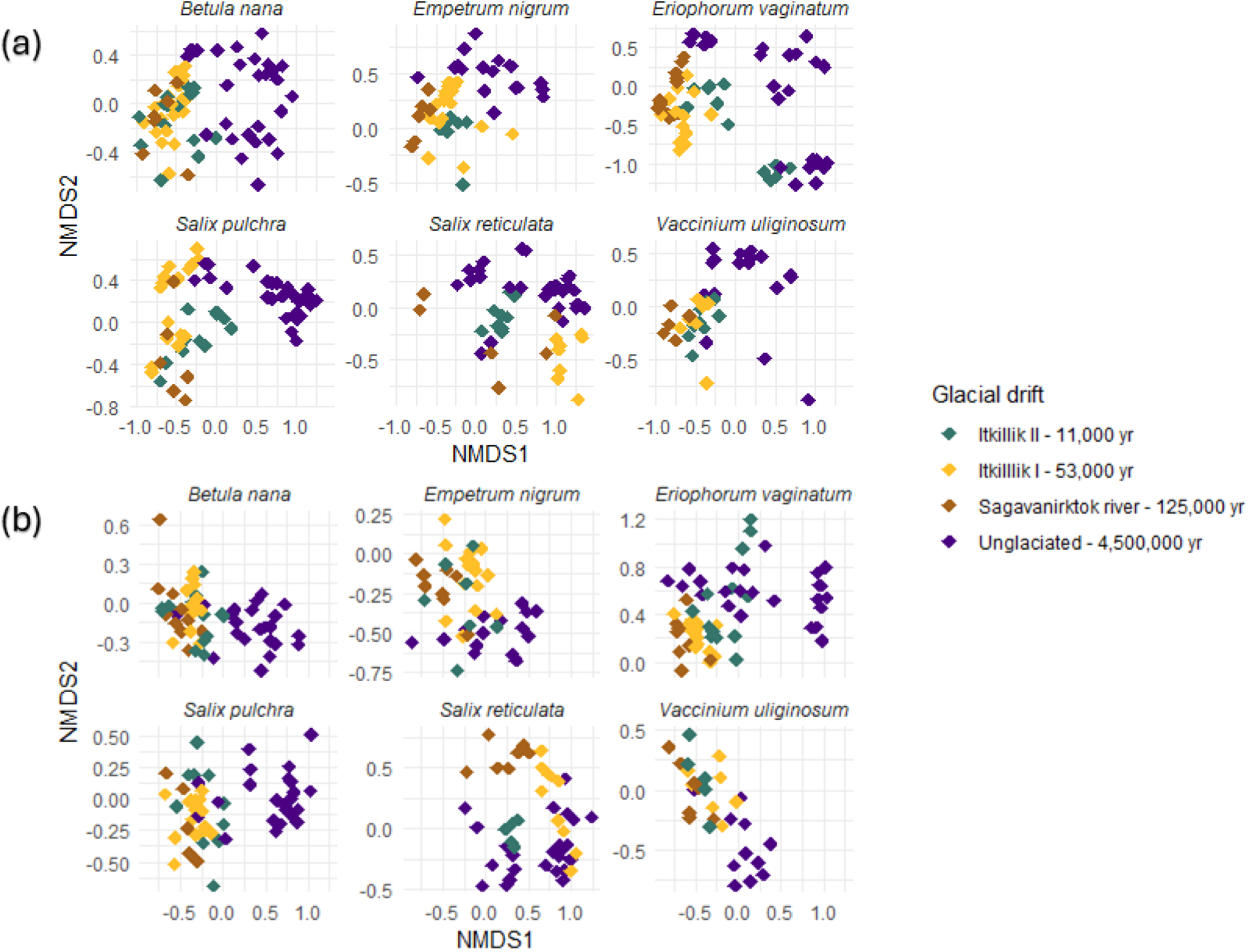
Rhizosphere bacteria (a) and fungi (b) group by glacial history across various host plant species. PERMANOVA results (Table 2) reveal significant groupings between glacial drifts (history) and host plant species which is visualized here. Ordinations were performed on the entire dataset and faceted by host plant species for visual clarity. Each point represents one rhizosphere community; colors represent the sites from which root samples were collected. Yr refers to the approximate number of years since the soil has been glaciated. The stress level was between 0.10 and 0.11 for 20 runs for bacterial ordinations and between 0.19 and 0.21 for 20 runs for fungal ordinations. Companion visualizations without facets can be found in (Fig. S6, S7).

### 3.3 ECM fungal communities vary most between sites and glacial drifts and vary least between plants at the same site or within roots of the same plant

When comparing ECM communities across glacial drifts, individual plants explained the most variation at 40.6% followed by glacial history at 10.2% (Table S2, Fig. S8, S9, S10). The variation of ECM communities *between sites* revealed individual plants explained 34.5% of variation while site explained 18.1% (Fig. 4, Table S2). To examine *within-site* variation we subset data by site and ran a series of PERMANOVA tests where individual plants were the only explanatory factor. We found that individual plants explained between 65% and 83% of *within-site* variation (Table S3). To examine within-plant variation, we chose 10 plants at random and compared the richness of ECM fungi across 3 root fragments from the same host plant (Table S4). Within an individual plant, species richness was variable, with some plants only containing 4 species of ECM fungi and others having as many as 60 species across all the root fragments (Fig. 3). The ECM fungal richness of a single *B. nana* root fragment ranged from 1 to 32 species with an average of 8.3 species per root fragment and a median of 7 species. A single root fragment was extremely variable in capturing ECM richness from an individual plant and ranged between 7% and 100%.

**Fig. 4.**
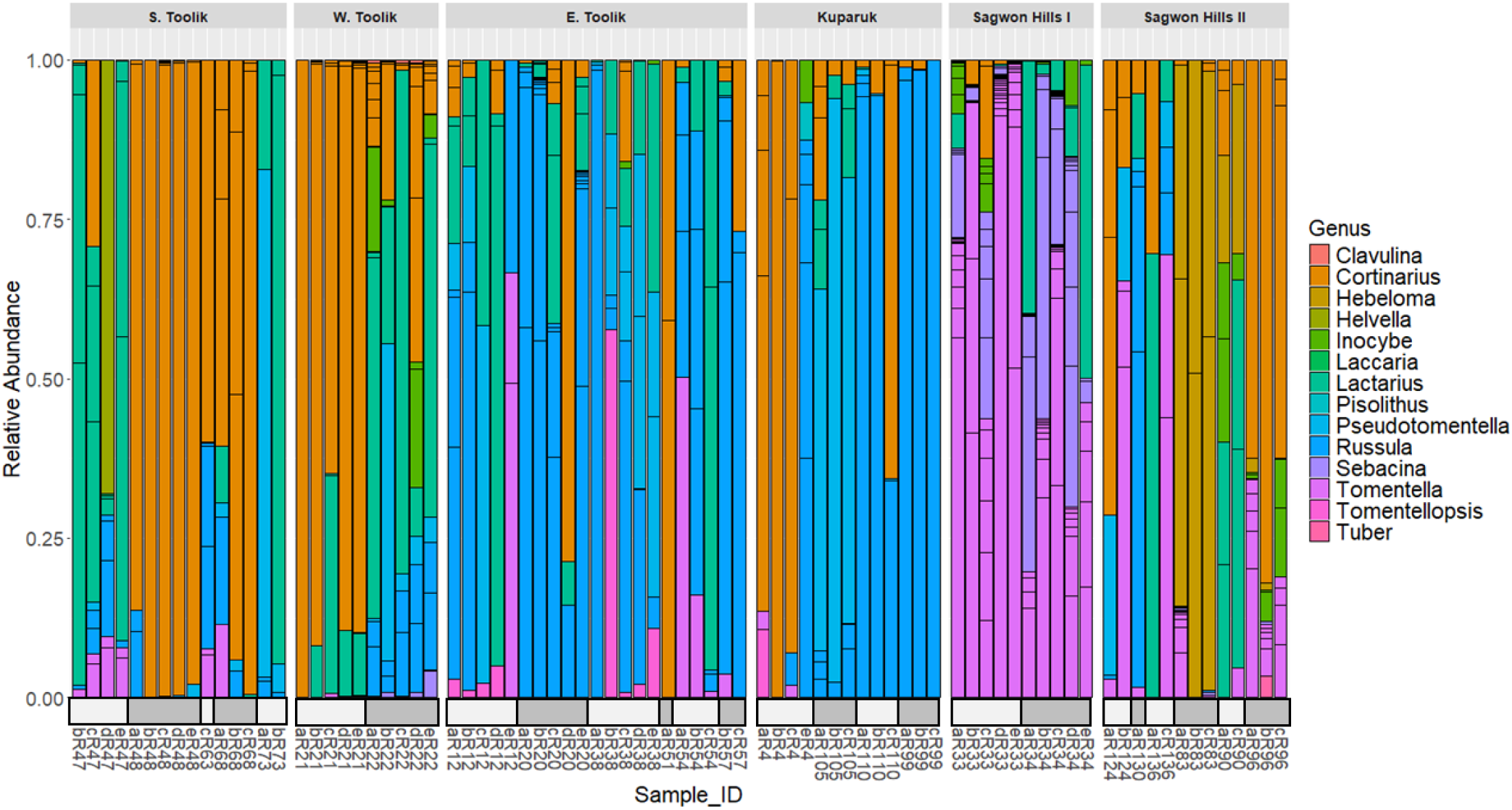
Sites and individual plants reveal distinct compositions of ECM fungal genera from *Betula nana* roots. Sample_ID represents a unique sample code where numbers refer to the individual plant and letters a-e represent the root fragment. Corresponding numbers have been grouped with white or gray boxes above the Sample_ID and represent fragments from the same plant (e.g., bR12, cR12, dR12). Plots are separated by sites and ASV counts were converted to relative abundance.

### 3.4 Ectomycorrhizae influences rhizosphere community composition

To determine the correlation between ECM fungi and rhizosphere communities we conducted Mantel tests comparing distance matrices of ECM fungi to rhizosphere bacteria, and ECM fungi to rhizosphere fungi. The composition of ECM fungal communities showed a significant correlation of 30.7% with bacterial composition of the rhizosphere (p < 0.001) (Fig. 5a) and a significant correlation of 54.7% of the fungal composition of the rhizosphere (p<0.001) (Fig. 5b).

**Fig. 5.**
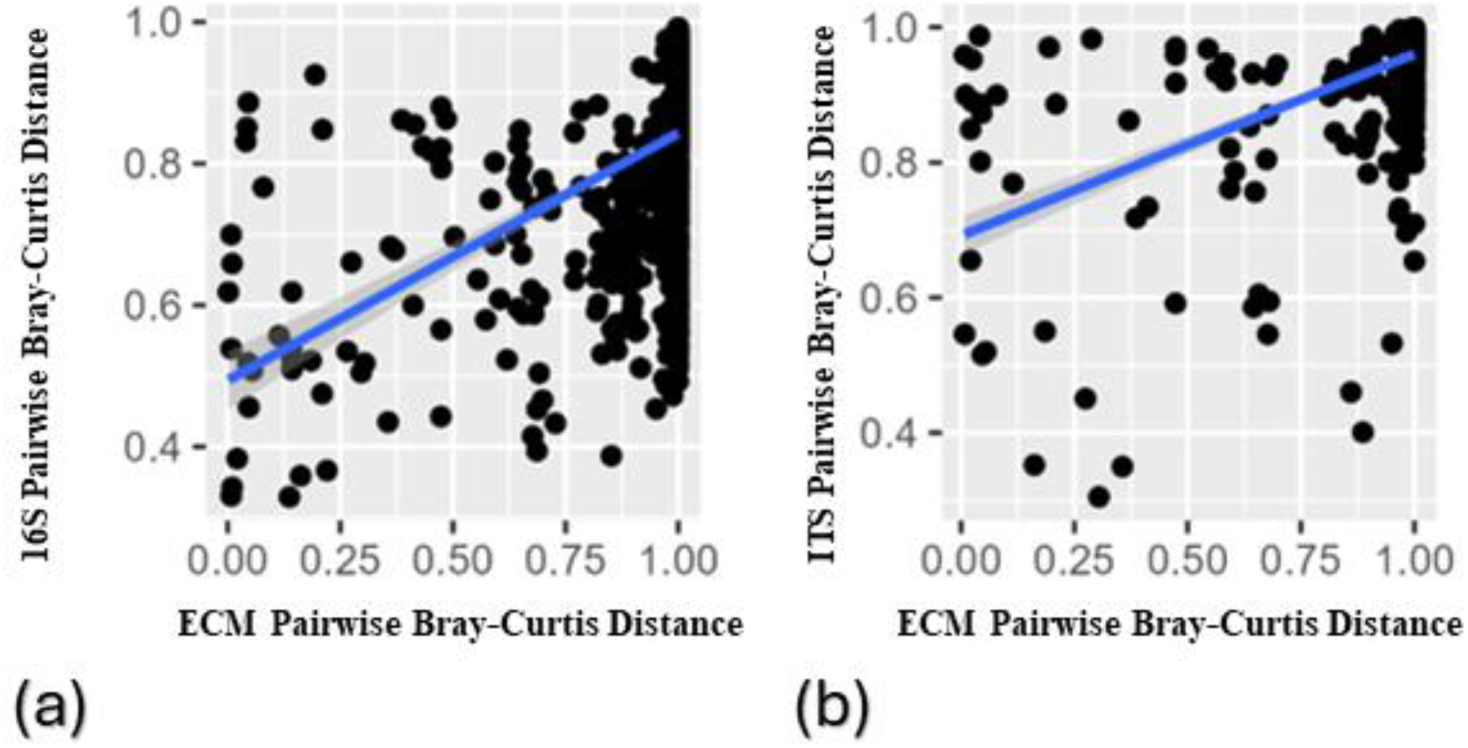
ECM fungi are highly correlated to rhizosphere bacterial and rhizosphere fungal communities. Mantel plots reveal high degrees of correlation between ECM fungal communities and **(a)** bacterial rhizosphere (Mantel R statistic = 0.31) as well as **(b)** fungal rhizosphere (Mantel R statistic = 0.55) communities suggesting co-occurrence of taxa across the respective communities. Each point corresponds to a root fragment where the Y-axis is a distance matrix generated from rhizosphere communities, and the X-axis is a distance matrix from ECM fungal communities. A trendline in blue depicts the Pearson correlation between the two species matrices. Self-comparisons have been removed from the plot.

### 3.5 Mycorrhizal type influences rhizosphere community composition

To explore how differences in mycorrhizal functional types (ie. ectomycorrhizal or ericoid fungi, or non-mycorrhizal) influence rhizosphere community composition we created an additional ordination based on mycorrhizal host (Fig. S11), as well as a separate PERMANOVA analysis for rhizosphere bacteria and fungi. We found that a plant’s mycorrhizal host type explained a significant amount of variation in rhizosphere communities accounting for 4.5% and 3.5% of variation in bacterial and fungal rhizosphere, respectively (Table S2). Since mycorrhizal type and host plant were confounded, and host plant explained more variation, we could not include mycorrhizal type in our main model.

## 4. Discussion

The physiology and metabolisms of tundra plants and their associated microbial communities are intimately intertwined (Fetcher and Shaver 1983, Shaver and Chapin 1991, Chapin et al., 1995, Hobbie and Chapin 1998, Weintraub and Schimel 2005, Wallenstein et al., 2007, Dunleavy and Mack 2021). Decades of research on plant species composition and physiology has led to the general understanding that plant root microbiomes are structured by plant genetics and environmental factors (Hartmann et al., 2009, Dennis et al., 2010, Liu et al., 2020, Liu et al., 2023). Here, we aim to deepen this understanding by exploring the relative contributions biotic and environmental factors play in structuring bacterial and fungal rhizosphere communities across different host plants and mineralogical soil types. We further explored the diversity of ECM fungi and their impact on rhizosphere microbiome structure in *B. nana*. We found the rhizosphere composition of tundra plants are shaped by glacial history, host plant species, vegetation community type, and mycorrhizal type, but that all tundra plants share a core microbiome (12.7% of bacteria and 5.5% of fungi). Likewise, we found that the most abundant 15 bacterial and fungal families comprise over 50% of the relative abundance of rhizosphere communities. As shrubification is rapidly shifting vegetation communities (Sturm 2001, Myers-Smith et al., 2011, Mekonnen et al., 2021), it is increasing the prevalence of obligate ECM and ERI fungi associated with shrubs (Gardes and Dahlberg 1996, Fujimara and Egger 2012, Clemmensen et al., 2015, Parker et al., 2015). Given that ECM and ERI fungi substantially impact microbial community composition and soil carbon processes in boreal systems (Averill and Hawkes 2016, Bödeker et al., 2016, Clemmensen et al., 2021), a better understanding of tundra rhizosphere community assemblages, and how they vary across host plants and soil types may enhance our ability to predict changes in microbial community composition and ecosystem processes brought on by shrubification. This work provides a foundation to better understand how changes in rhizosphere and mycorrhizal communities may be changing with shifting plant communities and soil types.

### 4.1 Tundra plants have a high amount of redundancy in rhizosphere community composition, yet the degree to which taxa are conserved differs dramatically between bacteria and fungi

There is a large amount of shared ASVs in rhizosphere communities for both bacteria and fungi. Interestingly, when we examine rhizosphere bacteria across host plants, we see that the largest class comparison consists of bacteria that are present in every host plant microbiome (12.7%), providing evidence that a substantial portion of the bacterial component of the rhizosphere of tundra plants is conserved independent of glacial history and plant species (Fig 1a). Likewise, we found a large degree of functional redundancy with little differences in the relative abundance of functional annotations in bacterial rhizosphere communities across host plants (Fig. S1) and glacial drifts (Fig. S2). We cannot rule out the fact that amplicon sequencing does not discriminate against living or non-living organisms, nor does it discriminate between active and dormant organisms; both of which may inflate the number of shared species across host plants (Burkert et al., 2019). Likewise, our functional predictions using FAPROTAX are bias towards culturable bacteria and may not accurately reflect whole communities *in situ* (Kuo et al., 2023). Nevertheless, we hypothesize that the high redundancy in bacterial composition and function across host plant microbiomes may be due to frequent disturbance events, such as cryoturbation and annual freeze thaw cycles, that select for bacteria capable of a wide range of metabolic parameters. For example, within intertidal systems, bacteria capable of metabolizing under various redox states were more abundant across the environment than more specialized bacteria (Chen et al., 2021). Because freeze-thaw cycles can alter moisture and redox potential in soil pore spaces (Rooney et al., 2022), metabolically flexible bacteria may have a competitive advantage that allows them to thrive in the rhizosphere of many host plants, especially when moisture or redox potentials change across seasons or topographical areas. A meaningful follow up study could be to assemble metagenomes from tundra rhizosphere communities and determine the proportion of genes that demonstrate high amounts of metabolic flexibility.

Although we did detect a core of shared fungal taxa present in all rhizosphere communities, the magnitude of shared fungi across host plant microbiomes was less pronounced than with bacteria, as 5.5% of fungal ASVs were shared across host plants compared to 12.7% for bacteria (Fig. 2b). We found that the five most abundant fungal class comparisons in our upset plots were ASVs specific to one plant rhizosphere and ranged from ⁓6.6% - 17.8% of ASVs. The proportion of ASVs specific to one host plant was higher for fungi than bacteria, suggesting more host specificity in rhizosphere fungi than bacteria. Additionally, across host plants we found 37% more fungal taxa were differentially abundant than bacteria (1,069 and 782, respectively; Fig. S4, SI Differential Abundance dataset), despite the observation that fungal richness was only one tenth of bacterial richness. Evidence for greater fungal host specificity relative to bacteria is also supported in temperate rhizosphere communities (Chen et al., 2022).

### 4.2 To what degree do glacial history and plant species explain rhizosphere community composition, and is this consistent for bacterial and fungal communities?

#### 4.2.1 Glacial history is a dominant driver of rhizosphere communities

Glacial history was the strongest factor explaining variation in bacterial rhizosphere communities and explained a significant amount of variation in fungal rhizosphere communities. Since glacial drift is a strong proxy for the stage of mineralogical weathering, it impacts many aspects of soil chemistry and biology (Schaetzl and Anderson 2006). Weathering of soil minerals influences edaphic factors including nutrient stabilization onto mineral surfaces (Doetterl et al., 2018), carbon destabilization (Bailey et al., 2019), and stimulation of SOM decomposition by plant inputs (Fang et al., 2023); as well as biological factors such as microbial community composition in the rhizosphere (Whitman et al., 2018) and bulk soils (Carson et al., 2007, Ahmed et al., 2017). Along our deglaciation gradient, we see stark differences in pH, %C and %N, extractable Fe and Al, and predominate mineral groups (Table 1). Using glacial history as a proxy for soil type, our results are consistent with prior studies that found soil type to be the strongest driver explaining rhizosphere composition in rice paddies (Liu et al., 2020a, Liu et al., 2020b), agricultural soils (Schlemper et al., 2018), and desert ecosystems (Muktar et al., 2021). Mineralogy and soil chemistry likely act as a first stage of environmental filtering in which microbes are preferentially selected based on their ability to live in environments with specific soil characteristics. This pool of environmentally filtered microbial communities in the bulk soil is then recruited to become part of the rhizosphere (de Vries and Wallenstein 2017). We observed this strong filtering by glacial history, despite our samples originating from the O-horizon; however, we did explain less variation than in previous studies that investigated deeper soil horizons (Liu et al., 2020a, Schlemper et al., 2018). This may be due to a reduction in the strength of the ecological filter imposed by mineralogy and soil properties without direct contact with minerals in the O-horizon. Nevertheless, the consistency of soil type as a primary driver of rhizosphere communities, in both mineral and organic horizons and across various ecosystems (including those in this study), highlights the importance of soil properties as an initial filter which then sets the stage for downstream metabolic and physiological processes which influence microbial community composition and biogeochemical processes.

#### 4.2.2 Host plant species and mycorrhizal types are significant drivers of rhizosphere composition

Rhizosphere communities are a fluid reflection of bulk soil communities, plant genetics, mycorrhizal type, plant developmental status, plant health, and nutrient foraging strategies (Jones et al., 2004. By altering the patterns of exudation—specifically the timing, composition, and amount of exudates—plants can stimulate or suppress the growth of specific microorganisms, enabling them to, at least in part, cultivate and tailor their microbiome to meet particular needs (Hartmann et al., 2009, Dennis et al., 2010, Jones et al., 2019). Although direct evidence is limited, mycorrhizal fungi are also thought to engage in exudation and shape the mycorrhizosphere in a manner similar to plants (Jones et al., 2004, Vik et al., 2013, Gorka et al., 2019, Frey, 2019). Our data supports the concept of a plant-induced microbiome specialization, as host plant species explained 8.2% variation in rhizosphere bacterial composition and 6.8% for rhizosphere fungi (Table 2), indicating plant species plays a key role in structuring the rhizosphere microbiome. These findings are similar to desert plant rhizosphere communities, where host plant species explained 9% of bacterial rhizosphere composition (Muktar et al., 2021).

Interestingly, we found that the interaction effects between glacial history and host plant species increased the explanatory power from 8.2% and 6.8% respectively, to 9.9% and 11.6% for rhizosphere bacteria and fungi. Taken together, this suggests that host plants belonging to a glacial history context have a heightened influence on structuring the rhizosphere community than either factor alone. This nuanced relationship between glacial history and host plants may be explained by a couple hypotheses. The first, and perhaps most obvious, is that variation in rhizosphere communities are attributed to differences in plant genotypes (Yu and Hochholdinger 2018, Zheng et al., 2023) and the interaction effects are inflated due to a high abundance of similar genotypes present in the population at a given site. A second hypothesis is that since mycorrhizal type was confounded with host plants, the heightened interaction effect between glacial drift and host plants is due to the different niches and microhabitats inhabited by ECM fungi (Bruns 1995). Mycorrhizal fungi exhibit clear patterns of distribution across the tundra (Fig. 4, S8, S9), and this patchiness is likely a product of mineralogical filtering and random drift (Rosling et al., 2003, Rosling et al., 2004, Zhang et al., 2023). We hypothesize that the mineralogical conditions of the soil, selects for mycorrhizal fungi suited to those environments, which in turn select for mycorrhizosphere members through their own exudation patterns. Parsing through the relative contributions between plant genotype and mycorrhizal composition may provide key insight as to understanding the mechanism by which rhizosphere communities are structured across the tundra and could explain why host plants, in the context of a particular glacial drift, explain more variation in rhizosphere communities than host plants alone.

#### 4.2.3 The composition of rhizosphere bacteria is more strongly influenced by glacial history than rhizosphere fungi

Microbial communities are assembled by co-occurring processes which are a product of a microbe’s fitness and adaptations, as well as dispersal, diversification, and demographic drift (Dini-Andreote et al., 2015). Environmental selection refers to any biotic or abiotic process in which the presence, absence, and relative abundance of a species is determined by ecological and deterministic factors (Vellend and Agrawal 2010). In soil ecology these can range in scale from climatic variables to plant species, soil mineralogy, and species interactions, which influence the survivability and reproductivity of a species. Stochasticity refers to processes where the presence, absence, or relative abundance of a species is due to chance and not a product of environmental selection or fitness and are associated with probabilistic dispersal and ecological drift (Chase and Myers 2011).

In our study, differences in the amount of variation attributed to glacial history for rhizosphere bacteria and fungi may be explained by the differences in which these communities are assembled. For instance, glacial history explains 13.3% of rhizosphere bacterial variation and 6.3% of fungi, indicating that soil mineralogy and edaphic factors have a stronger influence on rhizosphere bacteria than fungi. When using glacial history as a proxy for soil type, this concept is supported by previous research where soil type explains more variation in rhizosphere bacteria than fungi (Schlemper et al., 2018, Cui et al., 2019, Liu et al., 2020b). Stochastic assembly processes are stronger drivers of composition in both free-living and ECM fungal communities than deterministic processes. Dispersal limitation and drift (both of which are stochastic), are the strongest processes shaping fungal communities in grasslands and alpine environments (Guo et al., 2023, Zhang et al., 2023); whereas deterministic factors play a stronger role in structuring bacterial communities in tundra and alpine habitats (Doherty et al., 2020, Wang et al., 2024). The consistency between the patterns of our findings and prior findings—specifically that soil type explains more variation in rhizosphere bacteria than rhizosphere fungi—and previous work examining assembly processes in bacteria and fungi—specifically the lower contribution of deterministic selection in fungal assembly—suggests that mineralogical characteristics and associated differences in soil edaphic properties have a stronger role selecting rhizosphere bacterial composition than fungal composition, and that stochasticity plays a larger role is shaping the assembly of rhizosphere fungal communities.

### 4.3 What is the variation of ECM fungal composition between sites, between plants at the same site and within a single plant?

#### 4.3.1 ECM communities are structured strongly by site despite the abundance of potential hosts across the North Slope

Many ECM fungi are generalists able to form symbioses with many tundra shrubs (Gardes and Dahlberg 1996, Botnen et al., 2014), so we predicted that the within-site variation in composition would be roughly equal to the between-site variation in ECM fungal communities, and that every site would have similar community compositions (Fig. 3). We found that ECM fungal communities were strongly structured by site (18.1% of variation), and secondarily structured by glacial drift (10.2% of variation), although these variables were confounded (Table 2, Table S2). The stronger influence of site effects relative to glacial history on ECM fungal community composition could suggest a higher amount of stochasticity and dispersal limitation rather than deterministic factors, which is similar to previous findings (Zhang et al., 2023). Since pH has been shown to play a dominant role in shaping ECM fungal community composition (Carteron et al., 2020, Lang et al., 2021) it would stand to reason that the combination of glacial history and pH-driven vegetation communities would explain more ECM fungal community composition than one of those factors alone. However, we were unable to test this given the low representation of non-acidic vegetation samples in our dataset. It is also possible that ECM fungi are deterministically selected by another variable that is captured by site but is not related to glacial history, such as competition with other fungi, or predation by collembola (Kanters et al., 2015).

To determine within-site variation, we subset our data by site and ran PERMANOVA’s using individual plants as our factor and compared the relative variation explained by individual plants across the different site subsets. The effect of individual plants in the subset data ranged between 65% and 83% of the variation in ECM fungal communities (Table S3), compared to 40.6% of variation explained in the full ECM fungal PERMANOVA (Table S2). Unsurprisingly, when the data was subset by site, we found individual plants to explain a higher proportion of the variation in rhizosphere communities. This suggests that plants belonging to the same site have more similar ECM fungal communities to each other than plants at different sites and that between-site variation is greater than within-site variation for ECM fungal communities.

Within-plant variation of ECM fungal communities was assessed by calculating the species richness of a single fragment relative to the total species found in all the fragments from that plant (Table S4). We found that a single root fragment contained between 7% and 100% of all the ECM species found in that individual plant. On average, a single root fragment contained 42.3% of the total ECM from that plant indicating that ECM fungal richness and colonization on roots is highly variable and dependent on various factors such as resource partitioning and competition (Bruns 1995).

### 4.4 To what extent are rhizosphere and ECM fungal community compositions correlated, and does the strength of this correlation differ for ECM to bacterial and ECM to fungal rhizospheres?

#### 4.4.1 ECM fungal composition is significantly correlated to rhizosphere communities

The mycorrhizosphere is a hotbed for microbial activity, biomass, and biodiversity due to high rates of rhizodeposition and turnover of microbial necromass (Jones et al., 2004, Jones et al., 2009). Easily metabolized exudates, released from plant roots and mycorrhizal hyphae, fuel microbial growth and may offer a competitive advantage toward taxa that are better adapted for fast growth on those specific substrates (Marschmann et al., 2022). Root exclusion experiments have demonstrated ECM fungi engage in priming and C transfer (Brzostek et al., 2015, Gorka et al., 2019) and when combined with the observations that ECM fungi foster distinct microbial communities (Vik et al., 2013) and distinct exudate profiles from roots (Wang et al., 2017), we posit that ECM fungi influence soil chemistry and microbial communities by plant-independent mechanisms.

Our mantel test results revealed that ECM communities’ compositions are strongly correlated to both bacterial and fungal community compositions in the rhizosphere at 30.7% and 54.7% respectively (Fig. 4). The extent to which these relationships are causal or coincidental is unclear and literature suggests they may be highly context-dependent on the form and availability of N (Bhatnagar et al., 2018, Cui et al., 2019). In general, ECM fungi are better at accessing and acquiring organic forms of N and are typically associated with organic horizons, whereas saprotrophs with fresh leaf litter and mineral horizons (Lindahl et al., 2007, Bӧdeker et al., 2016). ECM fungi can both stimulate and suppress the growth of saprotrophic fungi (Gadgil and Gadgil 1971, Fernandez and Kennedy 2016); an ECM−saprotroph soil carbon model suggests that ECM fungi inhibit saprotrophs when litter inputs were relatively low in N, suggesting they have a stronger capacity to mine soil N than saprotrophs, and this suppression was mitigated when litter inputs were more labile (Shao et al., 2023). While it is impossible to fully disentangle the effect of the environment on both ECM fungal and rhizosphere communities, a possible explanation for the tight correlation between the two is that ECM and rhizosphere taxa are exerting some kind of biological filtering, or deterministic selection, on each other. ECM fungi may drive the selection of specific rhizosphere taxa through the production of exudates or metabolites, as ECM have distinct profiles of exudates and metabolites (Wang et al., 2017) which select for specific taxa when released into the mycorrhizosphere based on substrate use efficiencies and preference (Bhatnagar et al., 2018). Alternatively, an explanation could be that environmental factors are selecting for both ECM fungi and rhizosphere taxa, and the co-occurrence of taxa in the different functional types may be coincidence. For instance, edaphic factors such as moisture content, pH, substrate quality or accessibility to nutrients may select for particular members within ECM and rhizosphere groups, resulting in a high co-occurrence between taxa that prefer the same soil conditions. Substrates of a particular chemical composition, such as phenolics or those containing nutrients such as N or P, may select for ECM fungi and saprotrophs involved in the metabolism of that substrate (Talbot et al., 2013). Similarly, compounds found in micro-aggregates may select for filamentous fungi and yeasts able to access the compounds (Bach et al., 2018). Nevertheless, the strong relationship between ECM fungi and their rhizosphere counterparts suggests that microbial functions in the rhizosphere are either directly or indirectly influenced by the composition of ECM fungi.

## Supporting information

Differential Abundance Dataset

Relative Abundance Dataset

## Acknowledgments

We would like to thank Nathan Alexander and Ella Cotter for their help in processing rhizosphere and mycorrhizal samples and Dr. Eric Morrison for help in the DNA processing pipeline. We would like to thank Sarah Goldsmith, Else Schlermann and Amanda Young for helping in collection efforts at Toolik lake, Randy Fulweber for site selection, and Nate Blais, Thomas Muratore, and Josh Trombley for writing feedback. Finally, we would like to thank Dr. Stuart Grandy, Dr. Mark Waldrop and Dr. Ruth Varner for their contributions toward the experimental design.

## Author contributions

(CRediT authorship contribution statement)

**Sean Robert Schaefer:** Writing – original draft, Formal analysis, Investigation, Methodology, Project administration, Software, Visualization, Conceptualization, Data curation, Supervision.

**Fernando Montaño-Lopez**: Writing – review and editing, Data curation, Formal analysis. **Hannah Holland-Moritz**: Writing - review and editing, Software, Data curation, Conceptualization. **Caitlin Hicks Pries**: Writing – review and editing, Funding acquisition, Conceptualization, Resources. **Jessica Gilman Ernakovich**: Writing – review and editing, Funding acquisition, Conceptualization, Project administration, Resources, Supervision.

## Data availability

Sequence data is available on NCBI Sequence Read Archive: BioProject PRJNA1037070).

## References

Ahmed, E., Hugerth, L.W., Logue, J.B., Brüchert, V., Andersson, A.F., and Holmström, S. J. M. 2017. Mineral Type Structures Soil Microbial Communities. Geomicrobiology Journal 34: 538–545. doi:10.1080/01490451.2016.1225868.

Averill, C., and Hawkes, C.V. 2016. Ectomycorrhizal fungi slow soil carbon cycling. Ecology Letters 19: 937–947. doi:10.1111/ele.12631.

Bach, E.M., Williams, R.J., Hargreaves, S.K., Yang, F., and Hofmockel, K.S. 2018. Greatest soil microbial diversity found in micro-habitats. Soil Biology and Biochemistry 118: 217–226. 10.1016/j.soilbio.2017.12.018.

Badri, D.V., and Vivanco, J.M. 2009. Regulation and function of root exudates. Plant, cell & environment 32: 666–681. Wiley Online Library.

Bailey, V.L., Pries, C.H., and Lajtha, K. 2019. What do we know about soil carbon destabilization? Environmental Research Letters 14: 083004. doi:10.1088/1748-9326/ab2c11.

Barillot, C.D.C., Sarde, C.-O., Bert, V., Tarnaud, E., and Cochet, N. 2013. A standardized method for the sampling of rhizosphere and rhizoplan soil bacteria associated to a herbaceous root system. Annals of Microbiology 63: 471–476. doi:10.1007/s13213-012-0491-y.

Bell, C.W., Asao, S., Calderon, F., Wolk, B., and Wallenstein, M.D. 2015. Plant nitrogen uptake drives rhizosphere bacterial community assembly during plant growth. Soil Biology and Biochemistry 85: 170–182. 10.1016/j.soilbio.2015.03.006.

Bhatnagar, J.M., Peay, K.G., and Treseder, K.K. 2018. Litter chemistry influences decomposition through activity of specific microbial functional guilds. Ecological Monographs 88: 429–444. John Wiley & Sons, Ltd. doi:10.1002/ecm.1303.

Bödeker, I.T.M., Lindahl, B.D., Olson, Å., and Clemmensen, K.E. 2016. Mycorrhizal and saprotrophic fungal guilds compete for the same organic substrates but affect decomposition differently. Functional Ecology 30: 1967–1978. 10.1111/1365-2435.12677.

Botnen, S., Vik, U., Carlsen, T., Eidesen, P.B., Davey, M.L., and Kauserud, H. 2014. Low host specificity of root-associated fungi at an Arctic site. Molecular Ecology 23: 975–985. doi:10.1111/mec.12646.

Broeckling, C.D., Broz, A.K., Bergelson, J., Manter, D.K., and Vivanco, J.M. 2008. Root exudates regulate soil fungal community composition and diversity. Applied and environmental microbiology 74: 738–744. Am Soc Microbiol.

Bruns, T.D. 1995. Thoughts on the processes that maintain local species diversity of ectomycorrhizal fungi. Plant and Soil 170: 63–73. doi:10.1007/BF02183055.

Brzostek, E.R., Dragoni, D., Brown, Z.A., and Phillips, R.P. 2015. Mycorrhizal type determines the magnitude and direction of root-induced changes in decomposition in a temperate forest. New Phytologist 206: 1274–1282. 10.1111/nph.13303.

Burkert, A., Douglas, T.A., Waldrop, M.P., and Mackelprang, R. 2019. Changes in the Active, Dead, and Dormant Microbial Community Structure across a Pleistocene Permafrost Chronosequence. Applied and Environmental Microbiology 85: e02646–18. doi:10.1128/aem.02646-18.

Butler, B.M., and Hillier, S. 2021. powdR: An R package for quantitative mineralogy using full pattern summation of X-ray powder diffraction data. Computers & Geosciences 147: 104662. doi:10.1016/j.cageo.2020.104662.

Callahan, B.J., McMurdie, P.J., Rosen, M.J., Han, A.W., Johnson, A.J.A., and Holmes, S.P. 2016. DADA2: High-resolution sample inference from Illumina amplicon data. Nature Methods 13: 581–583. doi:10.1038/nmeth.3869.

Campbell, R., and Greaves, M.P. 1990. Anatomy and community structure of the rhizosphere. The rhizosphere.: 11–34. John Wiley and Sons Ltd.

Canarini, A., Kaiser, C., Merchant, A., Richter, A., and Wanek, W. 2019. Corrigendum: Root Exudation of Primary Metabolites: Mechanisms and Their Roles in Plant Responses to Environmental Stimuli. Frontiers in Plant Science 10. doi:10.3389/fpls.2019.00420.

Carson, J.K., Rooney, D., Gleeson, D.B., and Clipson, N. 2007. Altering the mineral composition of soil causes a shift in microbial community structure. FEMS Microbiology Ecology 61: 414–423. doi:10.1111/j.1574-6941.2007.00361.x.

Carter, M.R., and Gregorich, E.G. 2007. Soil Sampling and Methods of Analysis. CRC Press, Boca Raton. [Online] Available: 10.1201/9781420005271.

Carteron, A., Beigas, M., Joly, S., Turner, B.L., and Laliberté, E. 2020. Temperate Forests Dominated by Arbuscular or Ectomycorrhizal Fungi Are Characterized by Strong Shifts from Saprotrophic to Mycorrhizal Fungi with Increasing Soil Depth. Microb Ecol. doi:10.1007/s00248-020-01540-7.

Cazares, E. 1992. Mycorrhizal fungi and their relationship to plant succession in subalpine habitats. [Online] Available: https://api.semanticscholar.org/CorpusID:128216331.

Chapin, F.S., and Körner, Ch. 1995. Patterns, Causes, Changes, and Consequences of Biodiversity in Arctic and Alpine Ecosystems. Pages 313–320 in F.S. Chapin and C. Körner, eds. Arctic and Alpine Biodiversity: Patterns, Causes and Ecosystem Consequences. Springer Berlin Heidelberg, Berlin, Heidelberg. doi:10.1007/978-3-642-78966-3_22.

Chapin, F.S., Shaver, G.R., Giblin, A.E., Nadelhoffer, K.J., and Laundre, J.A. 1995. Responses of Arctic Tundra to Experimental and Observed Changes in Climate. Ecology 76: 694–711. Ecological Society of America. doi:10.2307/1939337.

Chase, J., and Myers, J. 2011. Chase JM, Myers JA.. Disentangling the importance of ecological niches from stochastic processes across scales. Philos Trans R Soc Lond B Biol Sci 366: 2351–2363. Philosophical transactions of the Royal Society of London. Series B, Biological sciences **366**: 2351–63. doi:10.1098/rstb.2011.0063.

Chen, Y., Xi, J., Xiao, M., Wang, S., Chen, W., Liu, F., Shao, Y., and Yuan, Z. 2022. Soil fungal communities show more specificity than bacteria for plant species composition in a temperate forest in China. BMC Microbiology 22: 208. doi:10.1186/s12866-022-02591-1.

Chen, Y.-J., Leung, P.M., Wood, J.L., Bay, S.K., Hugenholtz, P., Kessler, A.J., Shelley, G., Waite, D.W., Franks, A.E., Cook, P.L.M., and Greening, C. 2021. Metabolic flexibility allows bacterial habitat generalists to become dominant in a frequently disturbed ecosystem. ISME JOURNAL 15: 2986–3004. doi:10.1038/s41396-021-00988-w.

Cheng, W., and Coleman, D.C. 1990. Effect of living roots on soil organic matter decomposition. Soil Biology and Biochemistry 22: 781–787. 10.1016/0038-0717(90)90157-U.

Clemmensen, K.E., Durling, M.B., Michelsen, A., Hallin, S., Finlay, R.D., and Lindahl, B.D. 2021. A tipping point in carbon storage when forest expands into tundra is related to mycorrhizal recycling of nitrogen. Ecology Letters 24: 1193–1204. 10.1111/ele.13735.

Clemmensen, K.E., Finlay, R.D., Dahlberg, A., Stenlid, J., Wardle, D.A., and Lindahl, B.D. 2015.Carbon sequestration is related to mycorrhizal fungal community shifts during long-term succession in boreal forests. New Phytologist 205: 1525–1536. 10.1111/nph.13208.

Cui, Y., Bing, H., Fang, L., Wu, Y., Yu, J., Shen, G., Jiang, M., Wang, X., and Zhang, X. 2019. Diversity patterns of the rhizosphere and bulk soil microbial communities along an altitudinal gradient in an alpine ecosystem of the eastern Tibetan Plateau. Geoderma 338: 118–127. doi:10.1016/j.geoderma.2018.11.047.

Dennis, P.G., Miller, A.J., and Hirsch, P.R. 2010. Are root exudates more important than other sources of rhizodeposits in structuring rhizosphere bacterial communities? FEMS Microbiology Ecology 72: 313–327. doi:10.1111/j.1574-6941.2010.00860.x.

Doetterl, S., Berhe, A.A., Arnold, C., Bodé, S., Fiener, P., Finke, P., Fuchslueger, L., Griepentrog, M., Harden, J.W., Nadeu, E., Schnecker, J., Six, J., Trumbore, S., Van Oost, K., Vogel, C., and Boeckx, P. 2018. Links among warming, carbon and microbial dynamics mediated by soil mineral weathering. Nature Geoscience 11: 589–593. doi:10.1038/s41561-018-0168-7.

Doherty, S.J., Barbato, R.A., Grandy, A.S., Thomas, W.K., Monteux, S., Dorrepaal, E., Johansson, M., and Ernakovich, J.G. 2020. The Transition From Stochastic to Deterministic Bacterial Community Assembly During Permafrost Thaw Succession. Frontiers in Microbiology 11. doi:10.3389/fmicb.2020.596589.

Dunleavy, H.R., and Mack, M.C. 2021. Long-term experimental warming and fertilization have opposing effects on ectomycorrhizal root enzyme activity and fungal community composition in Arctic tundra. Soil Biology and Biochemistry 154: 108151. doi:10.1016/j.soilbio.2021.108151.

van Elsas, J.D., and Boersma, F.G.H. 2011. A review of molecular methods to study the microbiota of soil and the mycosphere. European Journal of Soil Biology 47: 77–87. 10.1016/j.ejsobi.2010.11.010.

Fang, Q., Lu, A., Hong, H., Kuzyakov, Y., Algeo, T.J., Zhao, L., Olshansky, Y., Moravec, B., Barrientes, D.M., and Chorover, J. 2023. Mineral weathering is linked to microbial priming in the critical zone. Nature Communications 14: 345. doi:10.1038/s41467-022-35671-x.

Fernandez, C.W., and Kennedy, P.G. 2016. Revisiting the ‘Gadgil effect’: do interguild fungal interactions control carbon cycling in forest soils? New Phytologist 209: 1382–1394. 10.1111/nph.13648.

Fetcher, N., and Shaver, G.R. 1983. Life Histories of Tillers of Eriophorum Vaginatum in Relation to Tundra Disturbance. Journal of Ecology 71: 131–147. [Wiley, British Ecological Society]. doi:10.2307/2259967.

Finley, B., Dijkstra, P., Rasmussen, C., Schwartz, E., Mau, R., Liu, X.-J.A., Van Gestel, N., and Hungate, B. 2018. Soil mineral assemblage and substrate quality effects on microbial priming. Geoderma 322: 38–47. doi:10.1016/j.geoderma.2018.01.039.

Finley, B.K., Mau, R.L., Hayer, M., Stone, B.W., Morrissey, E.M., Koch, B.J., Rasmussen, C., Dijkstra, P., Schwartz, E., and Hungate, B.A. 2021. Soil minerals affect taxon-specific bacterial growth. The ISME Journal. doi:10.1038/s41396-021-01162-y.

Fitch, A.A., Lang, A.K., Whalen, E.D., Geyer, K., and Hicks Pries, C. 2020. Fungal Community, Not Substrate Quality, Drives Soil Microbial Function in Northeastern U.S. Temperate Forests. Frontiers in Forests and Global Change 3. doi:10.3389/ffgc.2020.569945.

Fitch, A.A., Lang, A.K., Whalen, E.D., Helmers, E.M., Goldsmith, S.G., and Hicks Pries, C. 2023. Mycorrhiza Better Predict Soil Fungal Community Composition and Function than Aboveground Traits in Temperate Forest Ecosystems. Ecosystems. doi:10.1007/s10021-023-00840-6.

Frey, S.D. 2019. Mycorrhizal Fungi as Mediators of Soil Organic Matter Dynamics. Annual Review of Ecology, Evolution, and Systematics 50: 237–259. doi:10.1146/annurev-ecolsys-110617-062331.

von Fromm, S.F., Doetterl, S., Butler, B.M., Aynekulu, E., Berhe, A.A., Haefele, S.M., McGrath, S.P., Shepherd, K.D., Six, J., Tamene, L., Tondoh, E.J., Vågen, T.-G., Winowiecki, L.A., Trumbore, S.E., and Hoyt, A.M. 2024. Controls on timescales of soil organic carbon persistence across sub-Saharan Africa. Global Change Biology 30: e17089. John Wiley & Sons, Ltd. doi:10.1111/gcb.17089.

Fujimura, K.E., and Egger, K.N. 2012. Host plant and environment influence community assembly of High Arctic root-associated fungal communities. Fungal Ecology 5: 409–418. 10.1016/j.funeco.2011.12.010.

Gadgil, R.L., and Gadgil, P.D. 1971. Mycorrhiza and Litter Decomposition. Nature 233: 133–133. doi:10.1038/233133a0.

Gardes, M., and Dahlberg, A. 1996. Mycorrhizal diversity in arctic and alpine tundra: an open question. New Phytologist 133: 147–157. 10.1111/j.1469-8137.1996.tb04350.x.

Gorka, S., Dietrich, M., Mayerhofer, W., Gabriel, R., Wiesenbauer, J., Martin, V., Zheng, Q., Imai, B., Prommer, J., Weidinger, M., Schweiger, P., Eichorst, S.A., Wagner, M., Richter, A., Schintlmeister, A., Woebken, D., and Kaiser, C. 2019. Rapid Transfer of Plant Photosynthates to Soil Bacteria via Ectomycorrhizal Hyphae and Its Interaction With Nitrogen Availability. Front Microbiol 10: 168–168. doi:10.3389/fmicb.2019.00168.

Gough, L., Shaver, G., Carroll, J., Royer, D., and Laundre, J. 2000. Vascular plant species richness in Alaskan arctic tundra: The importance of soil pH. Journal of Ecology 88: 54–66. doi:10.1046/j.1365-2745.2000.00426.x.

Greaves, H.E., Eitel, J., Vierling, L., Boelman, N., Griffin, K., Magney, T., and Prager, C. 2019. High-Resolution Vegetation Community Maps, Toolik Lake Area, Alaska, 2013-2015. ORNL Distributed Active Archive Center. doi:10.3334/ORNLDAAC/1690.

Guo, Q., Wen, Z., Ghanizadeh, H., Fan, Y., Zheng, C., Yang, X., Yan, X., and Li, W. 2023. Stochastic processes dominate assembly of soil fungal community in grazing excluded grasslands in northwestern China. Journal of Soils and Sediments 23: 156–171. doi:10.1007/s11368-022-03315-8.

Hamilton, T.D. 2003. Glacial Geology of the Toolik Lake and Upper Kuparuk River Regions. University of Alaska. Institute of Arctic Biology. [Online] Available: http://hdl.handle.net/11122/1502.

Hartmann, A., Schmid, M., Tuinen, D. van, and Berg, G. 2009. Plant-driven selection of microbes. Plant and Soil 321: 235–257. doi:10.1007/s11104-008-9814-y.

Heijmans, M.M.P.D., Magnússon, R., Lara, M., Frost, G., Myers-Smith, I., Van Huissteden, K., Jorgenson, M., Fedorov, A., Epstein, H., Lawrence, D., and Limpens, J. 2022. Tundra vegetation change and impacts on permafrost. Nature Reviews Earth & Environment 3: 68–84. doi:10.1038/s43017-021-00233-0.

Hobbie, S., and Chapin, F. 1998. Response of tundra plant biomass, aboveground production, nitrogen, and CO2 flux to experimental warming. ECOLOGY 79: 1526–1544.

Iversen, C.M., Sloan, V.L., Sullivan, P.F., Euskirchen, E.S., McGuire, A.D., Norby, R.J., Walker, A.P., Warren, J.M., and Wullschleger, S.D. 2015. The unseen iceberg: plant roots in arctic tundra. New Phytologist 205: 34–58. 10.1111/nph.13003.

Jilling, A., Keiluweit, M., Contosta, A.R., Frey, S., Schimel, J., Schnecker, J., Smith, R.G., Tiemann, L., and Grandy, A.S. 2018. Minerals in the rhizosphere: overlooked mediators of soil nitrogen availability to plants and microbes. Biogeochemistry 139: 103–122. doi:10.1007/s10533-018-0459-5.

Jones, D.L., Hodge, A., and Kuzyakov, Y. 2004. Plant and mycorrhizal regulation of rhizodeposition. New Phytologist 163: 459–480. doi:10.1111/j.1469-8137.2004.01130.x.

Jones, D.L., Nguyen, C., and Finlay, R.D. 2009. Carbon flow in the rhizosphere: carbon trading at the soil–root interface. Plant and Soil 321: 5–33. doi:10.1007/s11104-009-9925-0.

Jones, P., Garcia, B.J., Furches, A., Tuskan, G.A., and Jacobson, D. 2019. Plant Host-Associated Mechanisms for Microbial Selection. Front Plant Sci 10: 862–862. doi:10.3389/fpls.2019.00862.

Kanters, C., Anderson, I., Johnson, D., Kanters, C., Anderson, I., and Johnson, D. 2015. Chewing up the Wood-Wide Web: Selective Grazing on Ectomycorrhizal Fungi by Collembola. FORESTS 6: 2560–2570. MDPI, Basel : doi:10.3390/f6082560.

Kluber, L.A., Smith, J.E., and Myrold, D.D. 2011. Distinctive fungal and bacterial communities are associated with mats formed by ectomycorrhizal fungi. Soil Biology and Biochemistry 43: 1042–1050. doi:10.1016/j.soilbio.2011.01.022.

Koizumi, T., and Nara, K. 2017. Communities of Putative Ericoid Mycorrhizal Fungi Isolated from Alpine Dwarf Shrubs in Japan: Effects of Host Identity and Microhabitat. Microbes Environ 32: 147–153. doi:10.1264/jsme2.ME16180.

Kolde, R. 2019. Pheatmap. [Online] Available: https://cran.r-project.org/package=pheatmap.

Kuo, J., Liu, D., and Lin, C.-H. 2023. Functional Prediction of Microbial Communities in Sediment Microbial Fuel Cells. Bioengineering 10. doi:10.3390/bioengineering10020199.

Lang, A.K., Jevon, F.V., Vietorisz, C.R., Ayres, M.P., and Hatala Matthes, J. 2021. Fine roots and mycorrhizal fungi accelerate leaf litter decomposition in a northern hardwood forest regardless of dominant tree mycorrhizal associations. New Phytologist 230: 316–326. 10.1111/nph.17155.

Lee, T., Walters, William A., Lennon, Niall J., Bochicchio, James, Krohn, Andrew, Caporaso, J. Gregory, and Pennanen, Taina 2016. Accurate Estimation of Fungal Diversity and Abundance through Improved Lineage-Specific Primers Optimized for Illumina Amplicon Sequencing. Applied and Environmental Microbiology 82: 7217–7226. American Society for Microbiology. doi:10.1128/AEM.02576-16.

Leopold, D.R. 2016. Ericoid fungal diversity: Challenges and opportunities for mycorrhizal research. Fungal Ecology 24: 114–123. doi:10.1016/j.funeco.2016.07.004.

Lindahl, B.D., Ihrmark, K., Boberg, J., Trumbore, S.E., Högberg, P., Stenlid, J., and Finlay, R.D. 2007. Spatial separation of litter decomposition and mycorrhizal nitrogen uptake in a boreal forest. New Phytol 173: 611–620. doi:10.1111/j.1469-8137.2006.01936.x.

Liu, L., Huang, X., Zhang, J., Cai, Z., Jiang, K., and Chang, Y. 2020a. Deciphering the relative importance of soil and plant traits on the development of rhizosphere microbial communities. Soil Biology and Biochemistry 148: 107909. 10.1016/j.soilbio.2020.107909.

Liu, L., Ma, L., Zhu, M., Liu, B., Liu, X., and Shi, Y. 2023. Rhizosphere microbial community assembly and association networks strongly differ based on vegetation type at a local environment scale. Frontiers in Microbiology 14. [Online] Available: https://www.frontiersin.org/journals/microbiology/articles/10.3389/fmicb.2023.1129471.

Liu, X.-J.A., Finley, B.K., Mau, R.L., Schwartz, E., Dijkstra, P., Bowker, M.A., and Hungate, B.A. 2020b. The soil priming effect: Consistent across ecosystems, elusive mechanisms. Soil Biology and Biochemistry 140: 107617. 10.1016/j.soilbio.2019.107617.

Loeppert, R., and Inskeep, W. 1996. Iron Methods of Soil Analysis. Part 3. Chemical Methods: 639– 664.

Louca, S., Parfrey, L.W., and Doebeli, M. 2016. Decoupling function and taxonomy in the global ocean microbiome. Science 353: 1272–1277. American Association for the Advancement of Science. doi:10.1126/science.aaf4507.

Love, M.I., Huber, W., and Anders, S. 2014. Moderated estimation of fold change and dispersion for RNA-seq data with DESeq2. Genome Biology 15: 550. doi:10.1186/s13059-014-0550-8.

Marschmann, G.L., Tang, J., Zhalnina, K., Karaoz, U., Cho, H., Le, B., Pett-Ridge, J., and Brodie, E.L. 2022. Life history strategies and niches of soil bacteria emerge from interacting thermodynamic, biophysical, and metabolic traits. bioRxiv: 2022.06.29.498137. doi:10.1101/2022.06.29.498137.

Marschner, P., Neumann, G., Kania, A., Weiskopf, L., and Lieberei, R. 2002. Spatial and temporal dynamics of the microbial community structure in the rhizosphere of cluster roots of white lupin (Lupinus albus L.). Plant and Soil 246: 167–174. doi:10.1023/A:1020663909890.

McMurdie, P. 2018.March 12. Functions for Accessing and (Pre)Processing Data. github. [Online] Available: https://joey711.github.io/phyloseq/preprocess.html [2022 Dec. 1].

McMurdie, P.J., and Holmes, S. 2013. phyloseq: An R package for reproducible interactive analysis and graphics of microbiome census data. PLoS ONE 8: e61217.

Mekonnen, Z.A., Riley, W.J., Berner, L.T., Bouskill, N.J., Torn, M.S., Iwahana, G., Breen, A.L., Myers-Smith, I.H., Criado, M.G., Liu, Y., Euskirchen, E.S., Goetz, S.J., Mack, M.C., and Grant, R.F. 2021. Arctic tundra shrubification: a review of mechanisms and impacts on ecosystem carbon balance. Environmental Research Letters 16: 053001. doi:10.1088/1748-9326/abf28b.

Miller, O.K. 1982. Mycorrhizae, Mycorrhizal Fungi and Fungal Biomass in Subalpine Tundra at Eagle Summit, Alaska. Holarctic Ecology 5: 125–134.

Molina, R., Massicotte, H., and Trappe, J. 1992. Specificity phenomena in mycorrhizal symbioses: Community-ecological consequences and practical implications. Mycorrhizal Functioning: An Integrative Plant-Fungal Process,.

Mukhtar, S., Mehnaz, S., and Malik, K.A. 2021. Comparative Study of the Rhizosphere and Root Endosphere Microbiomes of Cholistan Desert Plants. Frontiers in Microbiology 12. [Online] Available: https://www.frontiersin.org/articles/10.3389/fmicb.2021.618742.

Munroe, J.S., and Bockheim, J.G. 2001. Soil Development in Low-Arctic Tundra of the Northern Brooks Range, Alaska, U.S.A. Arctic, Antarctic, and Alpine Research 33: 78–87. Taylor & Francis. doi:10.1080/15230430.2001.12003407.

Muthukumar, T., Udaiyan, K., and Shanmughavel, P. 2004. Mycorrhiza in sedges—an overview. Mycorrhiza 14: 65–77. doi:10.1007/s00572-004-0296-3.

Myers-Smith, I.H., Forbes, B.C., Wilmking, M., Hallinger, M., Lantz, T., Blok, D., Tape, K.D., Macias-Fauria, M., Sass-Klaassen, U., Lévesque, E., Boudreau, S., Ropars, P., Hermanutz, L., Trant, A., Collier, L.S., Weijers, S., Rozema, J., Rayback, S.A., Schmidt, N.M., Schaepman-Strub, G., Wipf, S., Rixen, C., Ménard, C.B., Venn, S., Goetz, S., Andreu-Hayles, L., Elmendorf, S., Ravolainen, V., Welker, J., Grogan, P., Epstein, H.E., and Hik, D.S. 2011. Shrub expansion in tundra ecosystems: dynamics, impacts and research priorities. Environmental Research Letters 6: 045509. doi:10.1088/1748-9326/6/4/045509.

Nguyen, C. 2003. Rhizodeposition of organic C by plants: mechanisms and controls. Agronomie 23: 375–396.

Oksanen, J., Simpson, G.L., Blanchet, F.G., Kindt, R., Legendre, P., Minchin, P.R., O’Hara, R.B., Solymos, P., Stevens, M.H.H., Szoecs, E., Wagner, H., Barbour, M., Bedward, M., Bolker, B., Borcard, D., Carvalho, G., Chirico, M., Caceres, M.D., Durand, S., Evangelista, H.B.A., FitzJohn, R., Friendly, M., Furneaux, B., Hannigan, G., Hill, M.O., Lahti, L., McGlinn, D., Ouellette, M.-H., Cunha, E.R., Smith, T., Stier, A., Braak, C.J.F.T., and Weedon, J. 2022. vegan: Community Ecology Package. [Online] Available: https://CRAN.R-project.org/package=vegan.

Parada, A.E., Needham, D.M., and Fuhrman, J.A. 2016. Every base matters: assessing small subunit rRNA primers for marine microbiomes with mock communities, time series and global field samples. Environmental Microbiology 18: 1403–1414. John Wiley & Sons, Ltd. doi:10.1111/1462-2920.13023.

Parker, T.C., Subke, J.-A., and Wookey, P.A. 2015. Rapid carbon turnover beneath shrub and tree vegetation is associated with low soil carbon stocks at a subarctic treeline. Global Change Biology 21: 2070–2081. John Wiley & Sons, Ltd. doi:10.1111/gcb.12793.

Parker, T.C., Thurston, A.M., Raundrup, K., Subke, J.-A., Wookey, P.A., and Hartley, I.P. 2021. Shrub expansion in the Arctic may induce large-scale carbon losses due to changes in plant-soil interactions. Plant and Soil. doi:10.1007/s11104-021-04919-8.

Põlme, S., Abarenkov, K., Henrik Nilsson, R., Lindahl, B.D., Clemmensen, K.E., Havard, K., Nguyen, N., Kjøller, R., Bates, S.T., Baldrian, P., Frøslev, T.G., Adojaan, K., Vizzini, A., Suija, A., Pfister, D., Baral, H.-O., Järv, H., Madrid, H., Nordén, J., Liu, J.-K., Pawlowska, J., Põldmaa, K., Pärtel, K., Runnel, K., Hansen, K., Larsson, K.-H., Hyde, K.D., Sandoval-Denis, M., Smith, M.E., Toome-Heller, M., Wijayawardene, N.N., Menolli, N., Reynolds, N.K., Drenkhan, R., Maharachchikumbura, S.S.N., Gibertoni, T.B., Læssøe, T., Davis, W., Tokarev, Y., Corrales, A., Soares, A.M., Agan, A., Machado, A.R., Argüelles-Moyao, A., Detheridge, A., de Meiras-Ottoni, A., Verbeken, A., Dutta, A.K., Cui, B.-K., Pradeep, C.K., Marín, C., Stanton, D., Gohar, D., Wanasinghe, D.N., Otsing, E., Aslani, F., Griffith, G.W., Lumbsch, T.H., Grossart, H.-P., Masigol, H., Timling, I., Hiiesalu, I., Oja, J., Kupagme, J.Y., Geml, J., Alvarez-Manjarrez, J., Ilves, K., Loit, K., Adamson, K., Nara, K., Küngas, K., Rojas-Jimenez, K., Bitenieks, K., Irinyi, L., Nagy, L.G., Soonvald, L., Zhou, L.-W., Wagner, L., Aime, M.C., Öpik, M., Mujica, M.I., Metsoja, M., Ryberg, M., Vasar, M., Murata, M., Nelsen, M.P., Cleary, M., Samarakoon, M.C., Doilom, M., Bahram, M., Hagh-Doust, N., Dulya, O., Johnston, P., Kohout, P., Chen, Q., Tian, Q., Nandi, R., Amiri, R., Perera, R.H., dos Santos Chikowski, R., Mendes-Alvarenga, R.L., Garibay-Orijel, R., Gielen, R., Phookamsak, R., Jayawardena, R.S., Rahimlou, S., Karunarathna, S.C., Tibpromma, S., Brown, S.P., Sepp, S.-K., Mundra, S., Luo, Z.-H., Bose, T., Vahter, T., Netherway, T., Yang, T., May, T., Varga, T., Li, W., Coimbra, V.R.M., de Oliveira, V.R.T., de Lima, V.X., Mikryukov, V.S., Lu, Y., Matsuda, Y., Miyamoto, Y., Kõljalg, U., and Tedersoo, L. 2020. FungalTraits: a user-friendly traits database of fungi and fungus-like stramenopiles. Fungal Diversity 105: 1–16. doi:10.1007/s13225-020-00466-2.

R Core Team 2023. R: A Language and Environment for Statistical Computing. R Foundation for Statistical Computing, Vienna, Austria. [Online] Available: https://www.R-project.org/.

Rivers, A., Weber, K., Gardner, T., Liu, S., and Armstrong, S. 2018. ITSxpress: Software to rapidly trim internally transcribed spacer sequences with quality scores for marker gene analysis [version 1; peer review: 2 approved]. F1000Research **7**. doi:10.12688/f1000research.15704.1.

Rosling, A., Lindahl, B.D., Taylor, A.F.S., and Finlay, R.D. 2004. Mycelial growth and substrate acidification of ectomycorrhizal fungi in response to different minerals. FEMS Microbiology Ecology 47: 31–37. doi:10.1016/S0168-6496(03)00222-8.

Ryberg, M., Larsson, E., and Molau, U. 2009. Ectomycorrhizal Diversity on Dryas octopetala and Salix reticulata in an Alpine Cliff Ecosystem. Arctic Antarctic and Alpine Research - ARCT ANTARCT ALP RES 41: 506–514. doi:10.1657/1938-4246-41.4.506.

Schaetzl, R., and Anderson, S. 2006. Soil-Genesis and Geomorphology.

Schlemper, T.R., van Veen, J.A., and Kuramae, E.E. 2018. Co-Variation of Bacterial and Fungal Communities in Different Sorghum Cultivars and Growth Stages is Soil Dependent. Microbial Ecology 76: 205–214. doi:10.1007/s00248-017-1108-6.

Shao, S., Wurzburger, N., Sulman, B., and Hicks Pries, C. 2023. Ectomycorrhizal effects on decomposition are highly dependent on fungal traits, climate, and litter properties: A model-based assessment. Soil Biology and Biochemistry 184: 109073. doi:10.1016/j.soilbio.2023.109073.

Shaver, G.R., and Chapin, F.S. 1991. Production - Biomass Relationships and Element Cycling in Contrasting Arctic Vegetation Types. ECOLOGICAL MONOGRAPHS 61: 1–31. doi:10.2307/1942997.

Shirakawa, M., Uehara, I., and Tanaka, M. 2019. Mycorrhizosphere bacterial communities and their sensitivity to antibacterial activity of ectomycorrhizal fungi. Microbes and environments 34: 191–198. Japanese Society of Microbial Ecology/Japanese Society of Soil Microbiology ….

Smith, S.E., and Read, D.J. 2008. Mycorrhizal symbiosis. Academic press.

Smith-Schuyler, D. 2023. phylosmith: Functions to help analyze data as phyloseq objects. [Online] Available: https://schuyler-smith.github.io/phylosmith/.

Southard, J. 2022. 7.12: Glacial Deposits. Page in The Environment of the Earth’s Surface. Massachusetts Institue of Technology.

Sposito, G. 2008. The Chemistry of Soils. Oxford University Press.

Sturm, M., Racine, C., and Tape, K. 2001. Increasing shrub abundance in the Arctic. Nature 411: 546–547. doi:10.1038/35079180.

Talbot, J.M., Bruns, T.D., Smith, D.P., Branco, S., Glassman, S.I., Erlandson, S., Vilgalys, R., and Peay, K.G. 2013. Independent roles of ectomycorrhizal and saprotrophic communities in soil organic matter decomposition. Soil Biology and Biochemistry 57: 282–291. 10.1016/j.soilbio.2012.10.004.

Taudière, A. 2024. MiscMetabar: Miscellaneous Functions for Metabarcoding Analysis. [Online] Available: https://github.com/adrientaudiere/MiscMetabar.

Teixeira, C., Holland-Moritz, H., Granada, C., Bayer, C., Sausen, T., Tonial, F., Petry, C., and Frey, S. 2024. Land management of formerly subtropical Atlantic Forest reduces soil carbon stocks and alters microbial community structure and function. Applied Soil Ecology. doi:10.1016/j.apsoil.2023.105252.

Treu, R., Laursen, G.A., Stephenson, S.L., Landolt, J.C., and Densmore, R. 1995. Mycorrhizae from Denali National Park and Preserve, Alaska. Mycorrhiza 6: 21–29. doi:10.1007/s005720050101.

Vellend, M., and Agrawal, A. 2010. Conceptual Synthesis in Community Ecology. The Quarterly Review of Biology 85: 183–206. The University of Chicago Press. doi:10.1086/652373.

Vik, U., Logares, R., Blaalid, R., Halvorsen, R., Carlsen, T., Bakke, I., Kolstø, A.-B., Økstad, O.A., and Kauserud, H. 2013. Different bacterial communities in ectomycorrhizae and surrounding soil. Scientific Reports 3: 3471. doi:10.1038/srep03471.

de Vries, F.T., and Wallenstein, M.D. 2017. Below-ground connections underlying above-ground food production: a framework for optimising ecological connections in the rhizosphere. Journal of Ecology 105: 913–920. doi:10.1111/1365-2745.12783.

Walker, M.D., Walker, D.A., and Auerbach, N.A. 1994. Plant communities of a tussock tundra landscape in the Brooks Range Foothills, Alaska. Journal of Vegetation Science 5: 843–866. Wiley Online Library.

Wallenstein, M.D., McMahon, S., and Schimel, J. 2007. Bacterial and fungal community structure in Arctic tundra tussock and shrub soils. FEMS Microbiol Ecol 59: 428–35. doi:10.1111/j.1574-6941.2006.00260.x.

Wang, B., Chen, C., Xiao, Y.-M., Chen, K.-Y., Wang, J., Zhao, S., Liu, N., Li, J.-N., and Zhou, G.-Y. 2024. Trophic relationships between protists and bacteria and fungi drive the biogeography of rhizosphere soil microbial community and impact plant physiological and ecological functions. Microbiological Research 280: 127603. doi:10.1016/j.micres.2024.127603.

Wang, T., Tian, Z., Bengtson, P., Tunlid, A., and Persson, P. 2017. Mineral surface-reactive metabolites secreted during fungal decomposition contribute to the formation of soil organic matter. Environmental Microbiology 19: 5117–5129. John Wiley & Sons, Ltd. doi:10.1111/1462-2920.13990.

Weintraub, M.N., and Schimel, J.P. 2005. Nitrogen Cycling and the Spread of Shrubs Control Changes in the Carbon Balance of Arctic Tundra Ecosystems. BioScience 55: 408–415. doi:10.1641/0006-3568(2005)055[0408:Ncatso]2.0.Co;2.

Whitman, T., Neurath, R., Perera, A., Chu-Jacoby, I., Ning, D., Zhou, J., Nico, P., Pett-Ridge, J., and Firestone, M. 2018. Microbial community assembly differs across minerals in a rhizosphere microcosm. Environ Microbiol 20: 4444–4460. doi:10.1111/1462-2920.14366.

Whittinghill, K., and Hobbie, S. 2011. Effects of Landscape Age on Soil Organic MatterProcessing in Northern Alaska. Soil Science Society of America Journal 75: 907. doi:10.2136/sssaj2010.0318.

Wickham, H. 2016. ggplot2: Elegant Graphics for Data Analysis. Springer-Verlag New York. [Online] Available: https://ggplot2.tidyverse.org.

Wullschleger, S.D., Epstein, H.E., Box, E.O., Euskirchen, E.S., Goswami, S., Iversen, C.M., Kattge, J., Norby, R.J., van Bodegom, P.M., and Xu, X. 2014. Plant functional types in Earth system models: past experiences and future directions for application of dynamic vegetation models in high-latitude ecosystems. Annals of Botany 114: 1–16. doi:10.1093/aob/mcu077.

Yang, H., Zhao, X., Li, L., and Zhang, J. 2020. Detecting the colonization of ericoid mycorrhizal fungi in Vaccinium uliginosum using in situ polymerase chain reaction and green fluorescent protein. Plant Methods 16: 102. doi:10.1186/s13007-020-00645-x.

el Zahar Haichar, F., Santaella, C., Heulin, T., and Achouak, W. 2014. Root exudates mediated interactions belowground. Soil Biology and Biochemistry 77: 69–80. Elsevier.

Zhang, X., Wang, Y., Xu, Y., Babalola, B.J., Xiang, S., Ma, J., Su, Y., and Fan, Y. 2023. Stochastic processes dominate community assembly of ectomycorrhizal fungi associated with Picea crassifolia in the Helan Mountains, China. Frontiers in Microbiology 13. doi:10.3389/fmicb.2022.1061819.

