## Supplemental information for "Rhizosphere Bacteria and Fungi are Differentially Structured by Host Plants, Soil Mineralogy and Ectomycorrhizal Communities in the Alaskan Tundra"

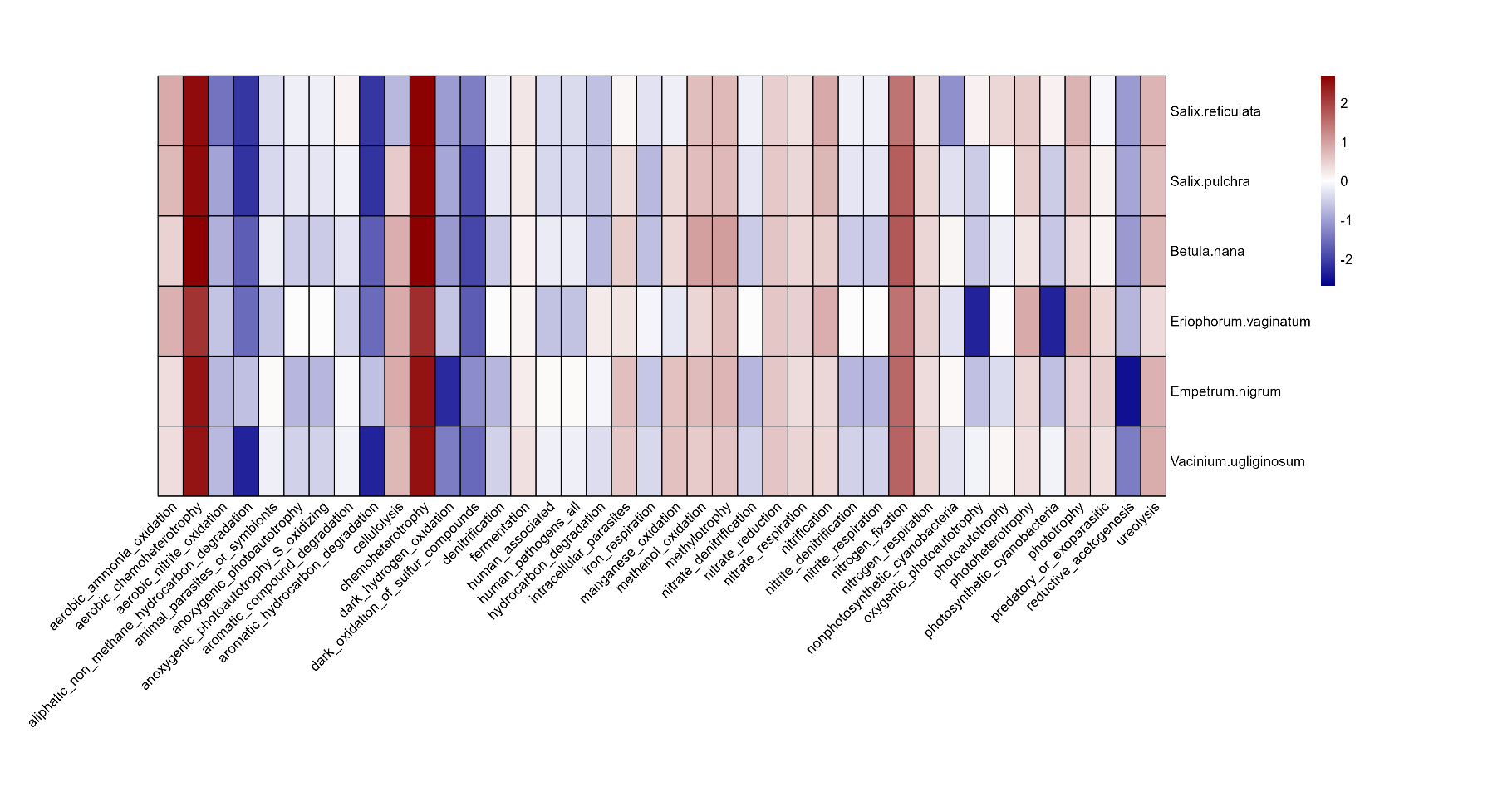

**Fig. S1. High counts of heterotrophic bacteria are present across host plants.** The mean number of ASVs for each host plant x functional annotation combination were obtained, log transformed, and relativized by plant species (rows). A total of 381 bacterial samples were annotated.

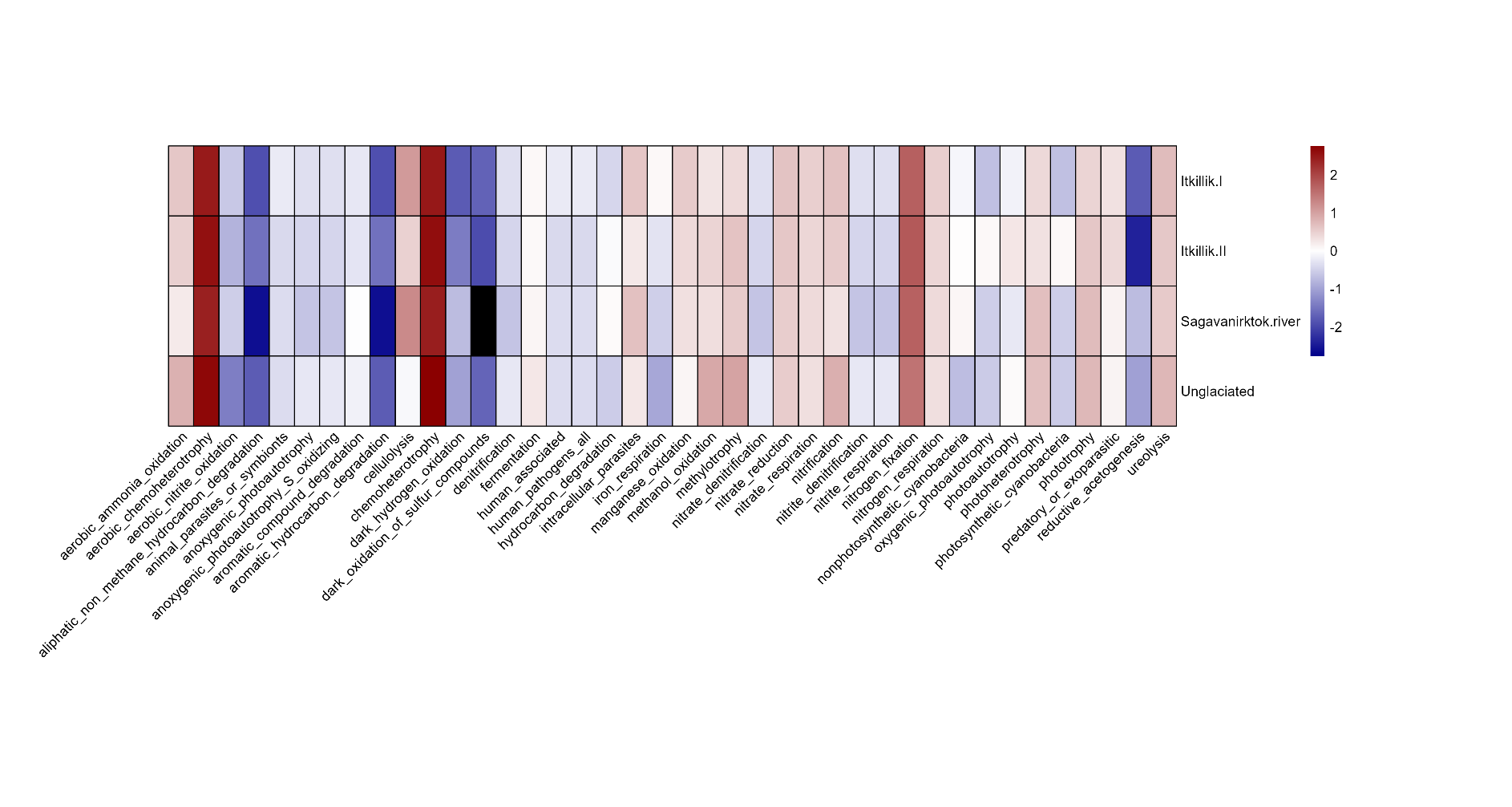

**Fig. S2. High counts of heterotrophic bacteria are present across glacial drifts.** The mean number of ASVs for each glacial drift x functional annotation combination were obtained, log transformed, and relativized by glacial drift (rows). Black filling denotes that no ASVs were detected. A total of 381 bacterial samples were annotated.

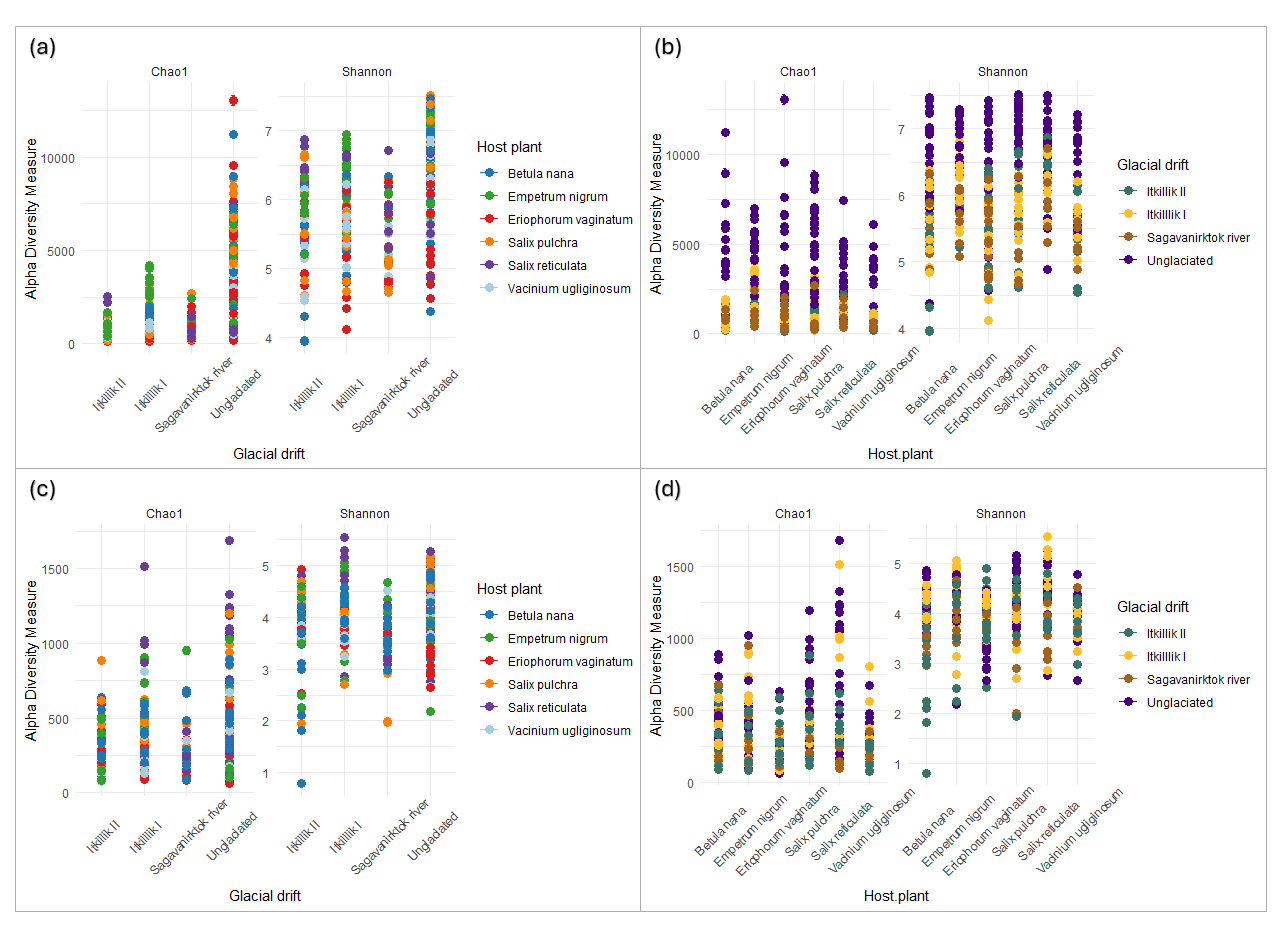

**Fig S3. Bacterial and Fungal Rhizosphere Diversity faceted by Glacial Drift and Host Plant.** Chao1 and Shannon diversity metrics indicate high levels of bacterial diversity **(a, b)** in *Empetrum nigrum* rhizospheres, as well as in Unglaciated sites; while Chao1 and Shannon diversity metrics indicate high levels of fungal diversity **(c, d)** in *Salix reticulata* and *Salix pulchra* rhizospheres, as well as in Unglaciated and Itkillik II sites, although this is less pronounced than bacterial diversity. Side by side comparisons are intended to better observe differences in diversity metrics across glacial histories (drift), host plants, and microbial kingdoms. Facets are separated by glacial drift **(a, c)** as well as host plant **(b, d)**.

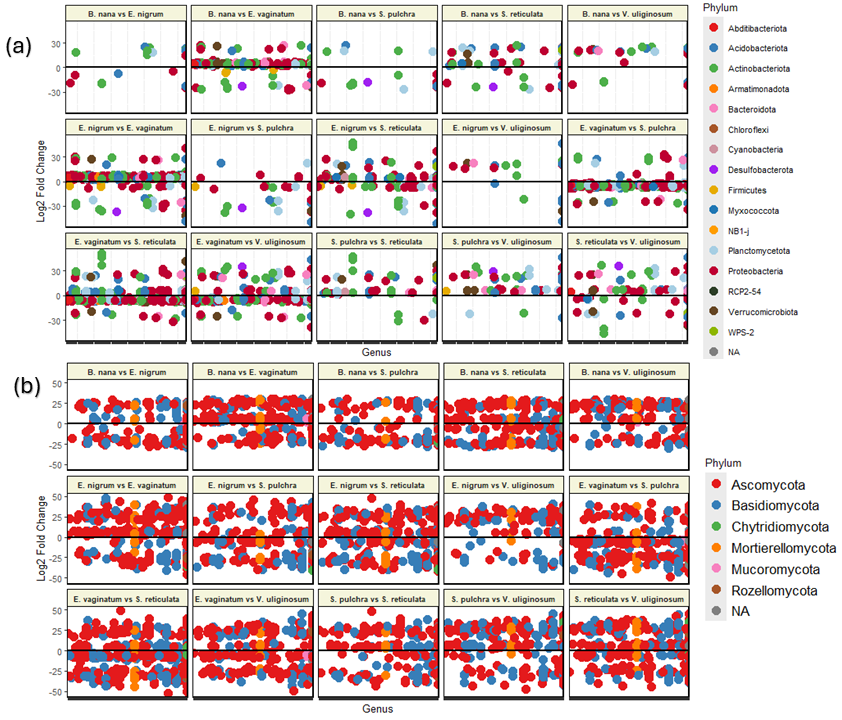

**Fig. S4. Plant species display differentially abundant microbial communities.** DASeq2 reveals 782 bacterial (a) and 1069 fungal (b) ASVs are significantly enriched or depleted relative to the plant they are being compared to. Color coding is relative to phylum level. Log2fold changes on the y-axis represent the magnitude of enrichment while the x-axis is organized alphabetically. Accompanying dataset found in supplemental files (Differential Abundance dataset).

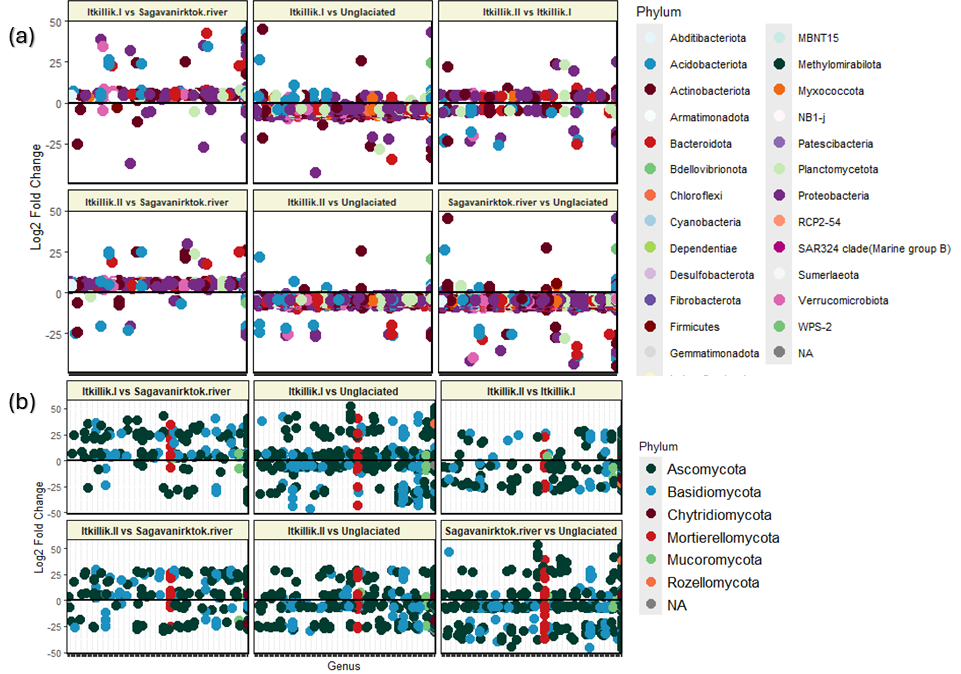

**Fig. S5. Glacial drifts display differentially abundant microbial communities.** DASeq2 reveals 2882 bacterial (a) and 769 fungal (b) ASVs are significantly enriched or depleted relative to the glacial drift they are being compared to. Color coding is relative to phylum level. Log2fold changes on the y-axis represent the magnitude of enrichment while the x-axis is organized alphabetically. Accompanying dataset found in supplemental files (Differential Abundance dataset).

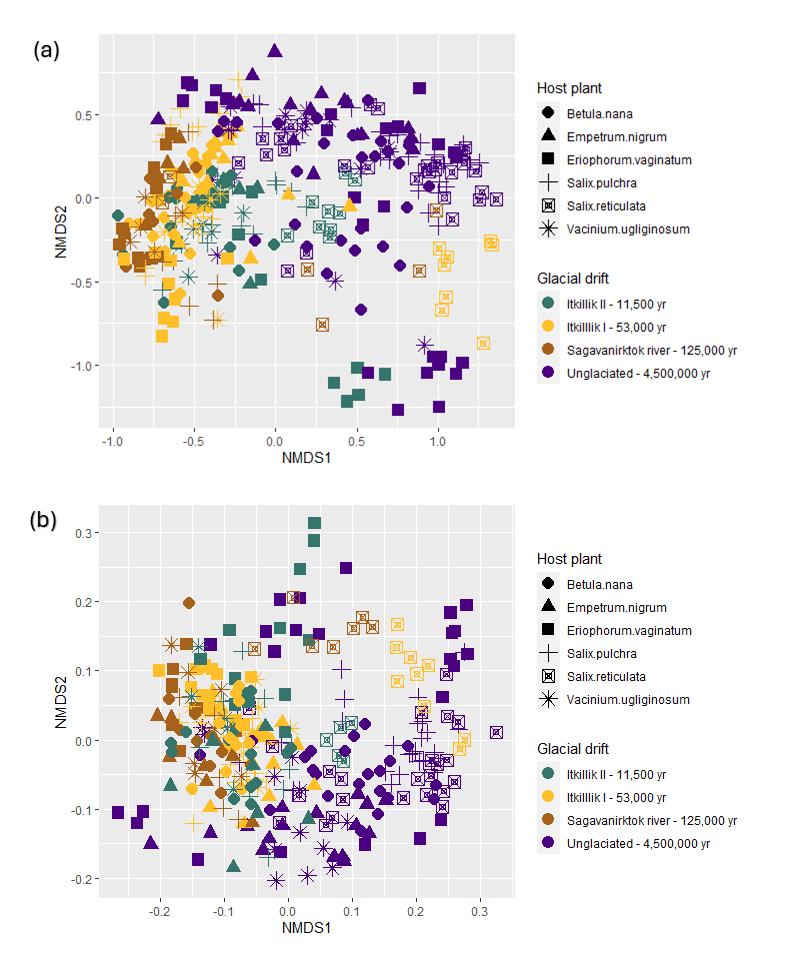

**Fig S6. Bacterial and Fungal rhizosphere NMDS ordination.** PERMANOVA results indicate significant groupings between host plants and glacial drifts (Table 2) for bacteria **(a)** and fungi **(b)**. Points represent rhizosphere communities; colors represent the glacial drifts from which root samples were collected from and shapes represent the host plant species rhizospheres were collected from. Yr refers to the approximate number of years since the soil has been glaciated. Bray-Curtis dissimilarity matrix was used. The stress level was between 0.10 and 0.11 for 20 runs for bacteria and between 0.19 and 0.21 for 20 runs for fungi.

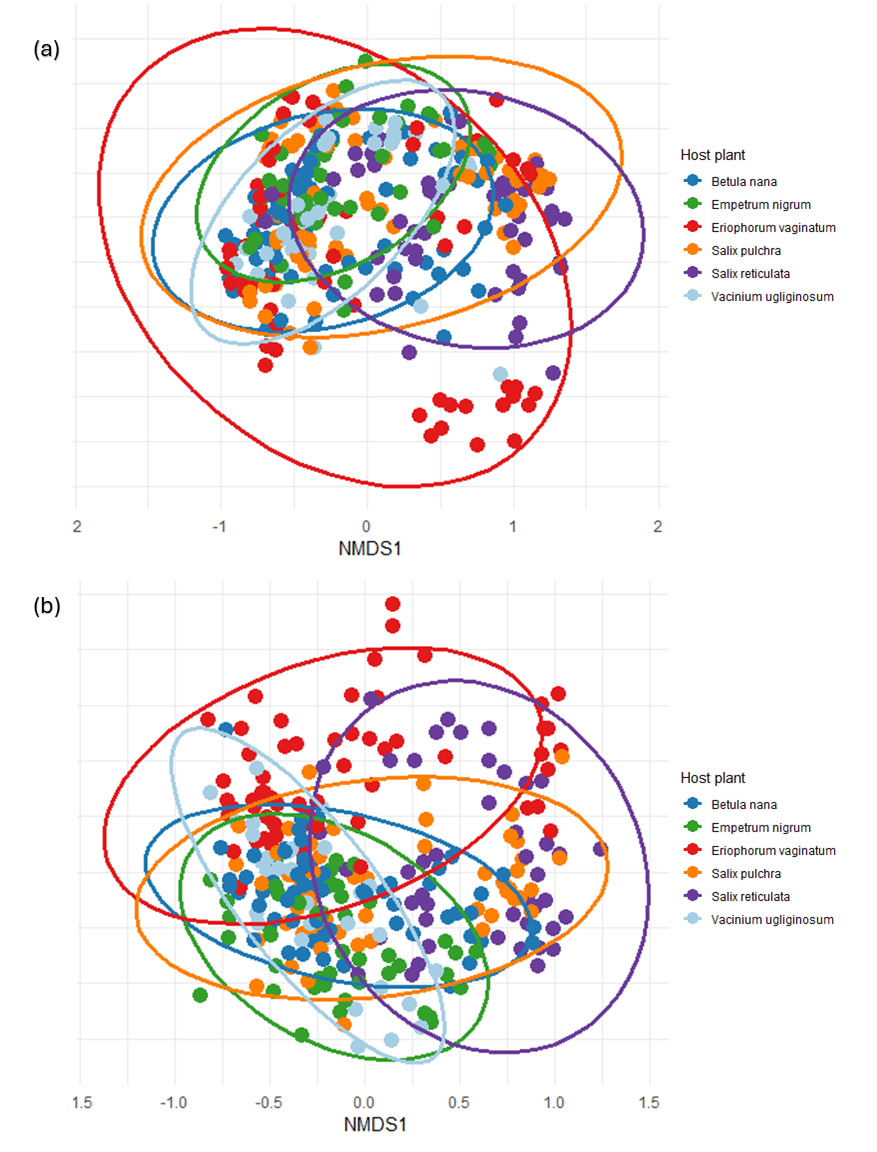

**Fig S7. Bacterial and fungal rhizosphere NMDS ordination colored by host plant.** PERMANOVA results indicate significant groupings between host plants (Table 2) for bacteria **(a)** and fungi **(b)**. Ellipses have been drawn to better differentiate between plants. Points represent rhizosphere communities; colors represent the glacial drifts from which root samples were collected from and shapes represent the host plant species rhizospheres were collected from. Bray-Curtis dissimilarity matrix was used. The stress level was between 0.10 and 0.11 for 20 runs for bacteria and between 0.19 and 0.21 for 20 runs for fungi.

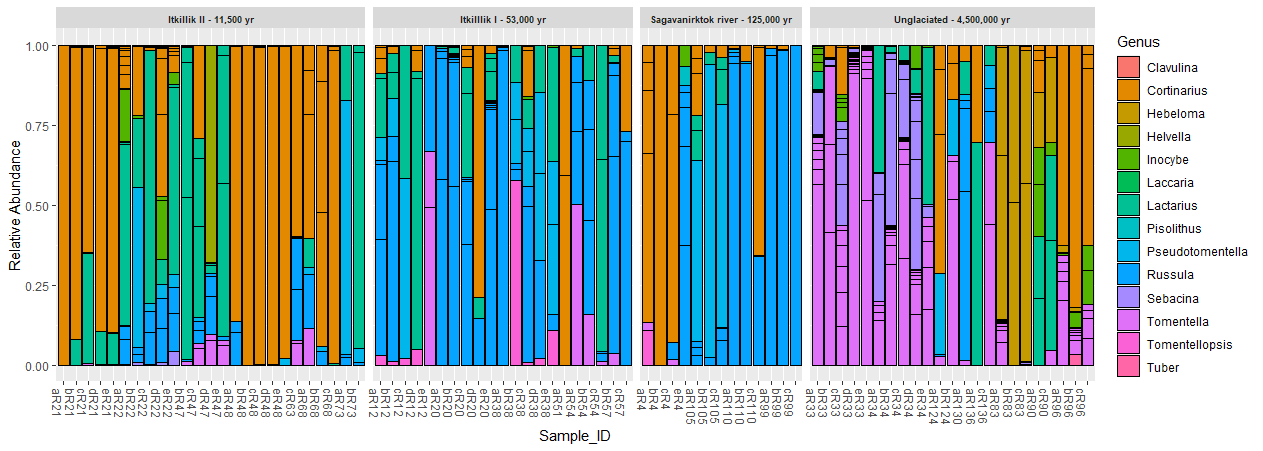

**Fig. S8. Glacial histories and individual plants reveal distinct compositions of ECM fungal genera from *Betula nana* roots.** Relative abundance counts from ECM fungal genera were obtained by normalizing sample reads and filtering for known ECM fungi in the Fungal Traits database. Sample_ID represents a unique sample code; numbers in the Sample_ID represent the individual plant and letters a-e represent the root fragment. Corresponding numbers represent fragments from the same plant (e.g., bR12, cR12, dR12). Plots are separated by glacial history.

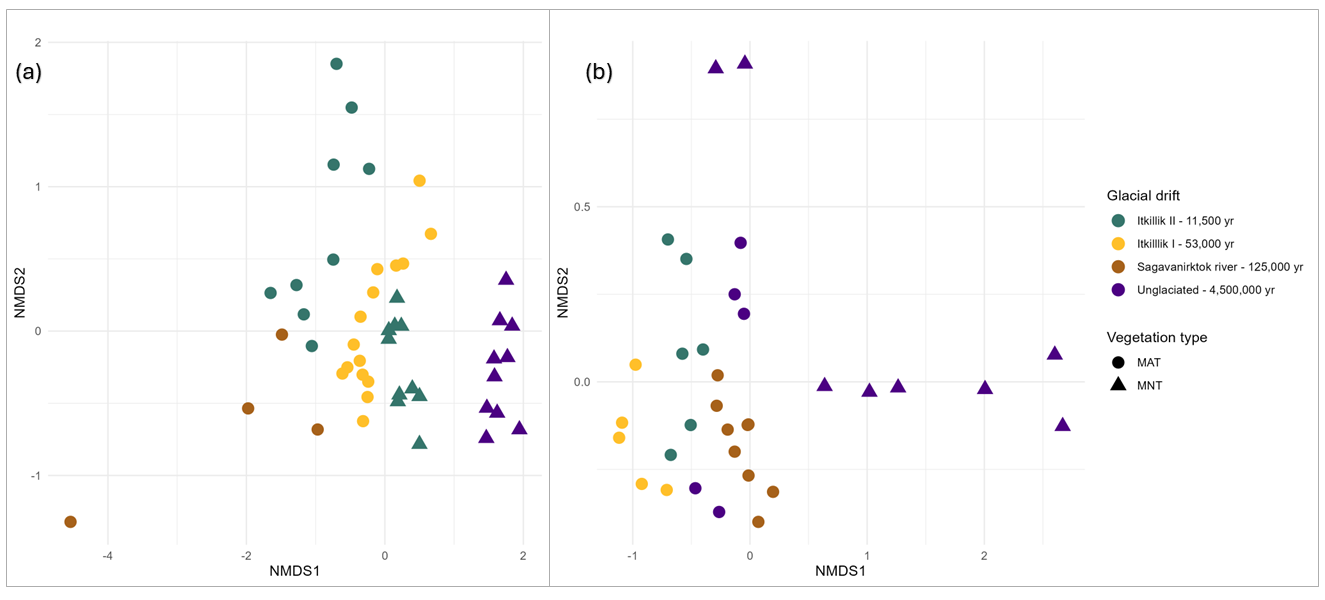

**Fig. S9. NMDS ordination of ECM fungal communities separated by glacial drifts and vegetation types.** To examine ECM community composition, we subset our fungal species matrix with only known ECM species reported in the Fungal Traits database. Due to the high level of heterogeneity between sample collection years including them on a single ordination was not informative. Instead, we chose to include two ordinations from the 2021 season **(a)** and 2022 season **(b)**. Points represent ECM communities on *B. nana* roots. Clustering of glacial history (drift) as well as vegetation communities can be visualized here. Yr refers to the approximate number of years since the soil has been glaciated. MAT refers to the vegetation communities, moist acidic tundra while MNT refers to moist non-acidic tundra. Bray-Curtis dissimilarity matrix was used and taxonomy was assigned to the genus level. The stress level was between 0.9 and 0.1 for the 2021 ordination **(a)** and between 0.06 and 0.07 for the 2022 ordination **(b)**. Both were for 20 runs.

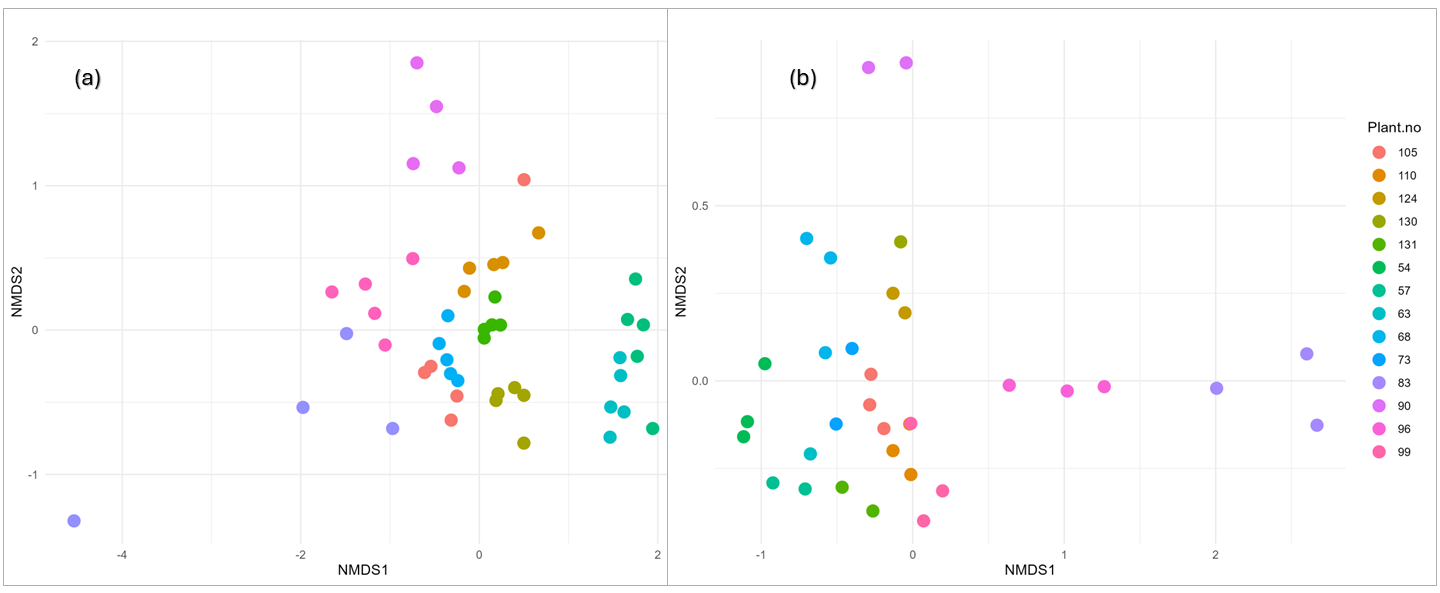

**Fig. S10. NMDS ordination of ECM fungal separated by individual plants.** To examine ECM community composition, we subset our fungal species matrix with only known ECM species reported in the Fungal Traits database. Due to the high level of heterogeneity between sample collection years including them on a single ordination was not informative. Instead, we chose to include two ordinations from the 2021 season **(a)** and 2022 season **(b)**. Points represent ECM communities on *B. nana* roots. Clustering of individual plants (plant. no) can be visualized here. Yr refers to the approximate number of years since the soil has been glaciated. MAT refers to the vegetation communities, moist acidic tundra while MNT refers to moist non-acidic tundra. Bray-Curtis dissimilarity matrix was used and taxonomy was assigned to the genus level. The stress level was between 0.9 and 0.1 for the 2021 ordination **(a)** and between 0.06 and 0.07 for the 2022 ordination **(b)**. Both were for 20 runs.

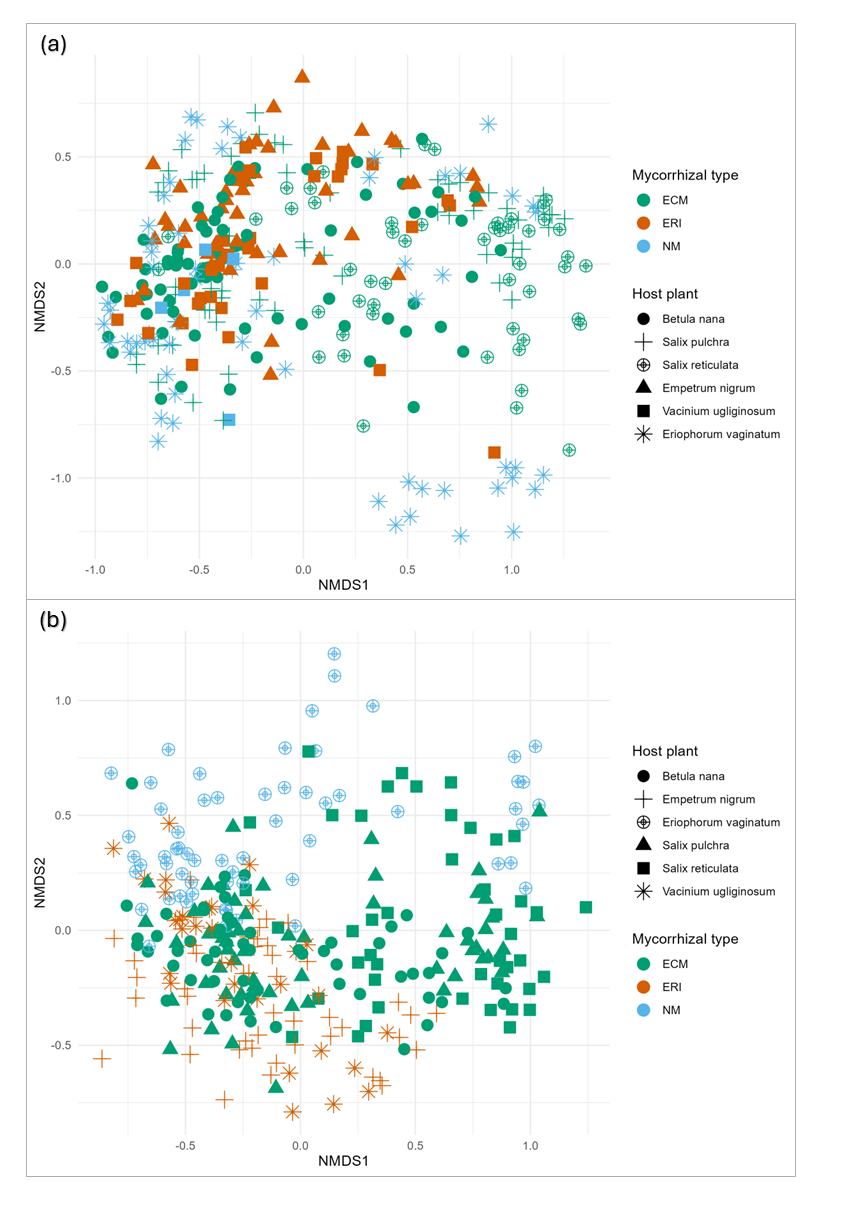

**Fig. S11. Bacterial and fungal rhizosphere communities group by mycorrhizal type.** PERMANOVA results indicate significant groupings between mycorrhizal host types (Table S4) for bacteria **(a)** and fungi **(b)**. Points represent rhizosphere communities; colors represent the glacial drifts from which root samples were collected from and shapes represent the host plant species rhizospheres were collected from. ECM refers to ectomycorrhizal fungi, ERI refers to ericoid mycorrhizae and NM is non-mycorrhizal. Bray-Curtis dissimilarity matrix was used and taxonomy was assigned to the genus level. The stress level was between 0.10 and 0.11 for 20 runs for both bacteria and fungi.

**Table S1. Samples used in rhizosphere and ECM fungal analyses.**

Due to PCR amplification efficiency and sequencing, as well as experimental design, the number of samples used in the various statistical tests varies. Each column represents a different statistical test used in this study. Non-metric dimensional scaling (NMDS) and PERMANOVA’s were conducted using the samples included in bacterial rhizosphere, fungal rhizosphere, and ECM fungal matrices. Mantel tests consisted of a subset of samples that were present in both the ECM fungal and rhizosphere matrices.

| **Glacial history** | **Site** | **Bacterial rhizosphere matrix (n)** | **Fungal rhizosphere matrix (n)** | **ECM fungal matrix (n)** | **Bacterial rhizosphere – ECM fungi mantel** | **Fungal rhizosphere – ECM mantel** |
| --- | --- | --- | --- | --- | --- | --- |
| **Itkillik II** | S. Dalton | 65 | 53 | 11 | 10 | 8 |
| **Itkillik II** | W. Toolik | 0 | 0 | 0 | 8 | 5 |
| **Itkillik I** | E. Toolik | 88 | 80 | 11 | 10 | 6 |
| **Sagavanirktok river (S.R.)** | Kuparuk | 44 | 46 | 9 | 9 | 9 |
| **Unglaciated** | Sagwon Hills I | 90 | 80 | 10 | 9 | 6 |
| **Unglaciated** | Sagwon Hills II | 54 | 43 | 6 | 6 | 5 |

**Table S2. PERMANOVA tables for bacterial and fungal models including mycorrhizal type instead of host plant, and ECM fungal model including site instead of glacial drift**

P values of <0.001 indicate significant groupings for all the factors tested in the model, suggesting they explain a high degree of variance in bacterial rhizosphere communities. Site refers to the operationally defined area from which we sampled from. Vegetation type refers to the vegetation communities, either moist acidic tundra (MAT) or moist non-acidic tundra (MNT). Individual plant refers to the plant which root fragments were separated from.

Glacial drift refers to the glacial history of the site and is a proxy for mineralogical weathering. Mycorrhizal type refers to the host plant’s mycorrhizal association and is either ectomycorrhizal, ericoid mycorrhizal or non-mycorrhizal.

| **Factor** | **Df** | **SumOfSqs** | **R2** | **F** | **Pr(>F)** |
| --- | --- | --- | --- | --- | --- |
| ***Bacterial model with mycorrhizal type instead of host plant*** | | | | | |
| Glacial drift | 3 | 15.02 | 0.133 | 18.67 | 0.001 |
| Mycorrhizal type | 2 | 5.058 | 0.045 | 9.427 | 0.001 |
| Vegetation type | 1 | 2.998 | 0.027 | 11.18 | 0.001 |
| Residual | 334 | 89.61 | 0.795 | - | - |
| Total | 340 | 112.7 | 1 | - | - |
| ***Fungal model with mycorrhizal type instead of host plant*** | | | | | |
| Glacial drift | 3 | 8.310 | 0.063 | 7.004 | 0.001 |
| Mycorrhizal type | 2 | 4.647 | 0.035 | 5.875 | 0.001 |
| Vegetation type | 1 | 2.712 | 0.020 | 6.857 | 0.001 |
| Residual | 295 | 116.7 | 0.882 | - | - |
| Total | 301 | 132.3 | 1 | - | - |
| ***ECM model with site instead of glacial drift (between sites)*** | | | | | |
| Site | 5 | 6.990 | 0.181 | 4.542 | 0.001 |
| Vegetation type | 1 | 0.792 | 0.021 | 2.575 | 0.001 |
| Individual plant | 17 | 13.31 | 0.345 | 2.544 | 0.001 |
| Residual | 57 | 17.54 | 0.454 | - | - |
| Total | 80 | 38.63 | 1 | - | - |

**Table S3. Compiled R^2^ values from site subsets for ECM fungal communities (within sites)**

To better examine within-site variation in ECM fungal communities, we ran linear models containing one site each and compiled the results here. P values of <0.001 indicate significant groupings for individual plants to explain variation in ectomycorrhizal communities. Because these are separate PERMANOVA results that have been compiled together, these results should be interpreted with caution and are intended to give a better sense of the range in which individual plants explain within-site variation of ECM communities. To this end, we subset our dataset by site and ran the linear model distance of species matrix ~ individual plant; permutations = 999. Because Sagwon Hills I and W. Toolik had only 2 individuals (10 subsamples each), we chose to not include them in this analysis as there was too few biological replicates.

| **Factor** | **Df** | **SumOfSqs** | **R2^*^** | **F** | **Pr(>F)** | **Site.name** |
| --- | --- | --- | --- | --- | --- | --- |
| Individual plant | 4 | 4.088 | **0.671** | 5.607 | 0.001 | Sagwon Hills II |
| Residual | 11 | 2.005 | 0.329 | NA | NA | Sagwon Hills II |
| Total | 15 | 6.093 | 1.000 | NA | NA | Sagwon Hills II |
| Individual plant | 4 | 2.840 | **0.705** | 3.590 | 0.001 | E.Toolik |
| Residual | 6 | 1.187 | 0.295 | NA | NA | E.Toolik |
| Total | 10 | 4.027 | 1.000 | NA | NA | E.Toolik |
| Individual plant | 4 | 3.324 | **0.833** | 7.476 | 0.002 | S.Dalton |
| Residual | 6 | 0.667 | 0.167 | NA | NA | S.Dalton |
| Total | 10 | 3.991 | 1.000 | NA | NA | S.Dalton |
| Individual plant | 3 | 1.871 | **0.651** | 3.111 | 0.013 | Kuparuk |
| Residual | 5 | 1.002 | 0.349 | NA | NA | Kuparuk |
| Total | 8 | 2.873 | 1.000 | NA | NA | Kuparuk |

**Table S4. ECM species richness for individual root fragments**

Because root subsamples are inherently pseudo-replicates and not true biological replicates, we could not run a linear model examining the amount of variation explained by a given root subsample. Instead, to get a sense for how much variation was captured by a root fragment (ie. aR, bR, cR…etc), we randomly chose 10 individual plants (denoted by the numbers, ie. 12, 20, 21…etc) and summed up the number of ECM fungal species present on all of the root fragments. We then deduced the proportion of ECM fungi captured on a given root fragment by dividing the number of ECM fungi found of that fragment divided by the total number of ECM fungi found in all of the root fragments from that individual plant. We then took an average of the proportion of species captured by each root fragment for an individual and found that on average, any given root fragment captured between 23% and 74%. The range in which any single root fragment captured ECM fungal communities was between 7% and 100%.

| Root fragment ID | Richness | Total species present | Proportion captured | Average proportion captured |
| --- | --- | --- | --- | --- |
| aR_12 | 10 | 15 | 0.67 | 0.47 |
| bR_12 | 8 | 15 | 0.53 |  |
| cR_12 | 3 | 15 | 0.20 |  |
| aR_20 | 6 | 22 | 0.27 | 0.44 |
| bR_20 | 14 | 22 | 0.64 |  |
| cR_20 | 9 | 22 | 0.41 |  |
| aR_21 | 2 | 28 | 0.07 | 0.23 |
| bR_21 | 10 | 28 | 0.36 |  |
| cR_21 | 7 | 28 | 0.25 |  |
| aR_33 | 25 | 60 | 0.42 | 0.31 |
| bR_33 | 14 | 60 | 0.23 |  |
| cR_33 | 17 | 60 | 0.28 |  |
| aR_34 | 12 | 57 | 0.21 | 0.36 |
| bR_34 | 18 | 57 | 0.32 |  |
| cR_34 | 32 | 57 | 0.56 |  |
| aR_68 | 7 | 11 | 0.64 | 0.52 |
| bR_68 | 5 | 11 | 0.45 |  |
| cR_68 | 5 | 11 | 0.45 |  |
| aR_96 | 10 | 21 | 0.48 | 0.52 |
| bR_96 | 11 | 21 | 0.52 |  |
| cR_96 | 12 | 21 | 0.57 |  |
| aR_99 | 3 | 4 | 0.75 | 0.58 |
| bR_99 | 3 | 4 | 0.75 |  |
| cR_99 | 1 | 4 | 0.25 |  |
| aR_105 | 9 | 9 | 1.00 | 0.74 |
| bR_105 | 4 | 9 | 0.44 |  |
| cR_105 | 7 | 9 | 0.78 |  |
| aR_110 | 7 | 8 | 0.88 | 0.58 |
| bR_110 | 3 | 8 | 0.38 |  |
| cR_110 | 4 | 8 | 0.50 |  |
